# Social connections, group size and fine-scale manipulations of social stability shape learning, scrounging and foraging rate in mixed flocks of wild songbirds

**DOI:** 10.1101/2025.08.18.670876

**Authors:** Michael S. Reichert, Josh A. Firth, Camille Troisi, John L. Quinn

**Affiliations:** Department of Biology, Oklahoma State University, Stillwater, Oklahoma, USA; School of Biological, Earth and Environmental Sciences, University College Cork, Cork, Ireland; Department of Biology, Edward Grey Institute, University of Oxford, Oxford, U.K.; Faculty of Biological Sciences, University of Leeds, Leeds, U.K.; Ethologie animale et humaine, Université de Rennes, Rennes, France

**Keywords:** social foraging, production learning, scrounging, RFID, group size, foraging strategy

## Abstract

The social environment provides both opportunities and challenges for foragers. How multiple components of sociality and foraging interact simultaneously is poorly understood, while experimental manipulation of sociality within wild foraging groups is rare. We tested whether production learning, scrounging, and foraging rate are associated with individual social network metrics and manipulated social stability in mixed species flocks of wild birds. First, individuals were randomly allowed access to, and learned to forage primarily from, one of five feeders. Thereafter, we performed two reversals, manipulating social connections by either assigning birds to a new feeder along with all others assigned to their previous feeder (stable treatment) or reassigning each bird individually (unstable treatment). Most of the effects observed were context dependent but in general we found that: i) Learning was slower in the stable treatment, for individuals with higher weighted degree, and in smaller flocks; ii) Scrounging was higher in the stable treatment, increased with weighted degree, but decreased with flock size; and iii) Foraging rate was predicted only by weighted degree. Our findings demonstrate the complex and dynamic nature of the relationship between sociality and foraging and suggest that selection for group living is more nuanced than usually recognized.

## INTRODUCTION

Foraging is the result of a complex set of decisions about when, where and how to obtain food (Stevens et al. 2007). Because foraging provides the energy that is critical for fitness, a diverse set of strategies have evolved to ensure foraging success (Waite and Field 2007). A key challenge for foragers is that this behavior takes place in a constantly changing environment, and this environmental change can drastically affect variables that determine the risks and rewards of any given foraging strategy such as the availability and distribution of food, the technique required to extract it, and the necessary trade-offs with other traits such as reproduction and vigilance against predators (Oom et al. 2004; Monk et al. 2018).

Additionally, for the many species that forage in social groups, sociality can offer both opportunities and challenges to foraging individuals (Clark and Mangel 1986; Krause and Ruxton 2002). Social groups may provide advantages to individual foragers such as increased discovery of food (Pitcher et al. 1982; Snijders et al. 2021), increased efficiency in extracting it (Courchamp and Macdonald 2001; Beauchamp 2005; MacNulty et al. 2014), opportunities to socially learn from other individuals (Hoppitt and Laland 2013) and shared vigilance against predators (Cresswell 1994; Lima et al. 1999; Silk 2007; Sansom et al. 2008). However, foraging in a group also leads to increased competition (Cresswell 1997; Focardi and Pecchioli 2005; Beltrão et al. 2022), as well as other general costs of sociality such as predator attraction and disease spread (Godfrey et al. 2009; Drewe 2010; Weber et al. 2013). Previous studies show mixed results for whether social group characteristics such as group size enhance or detract from foraging success (Beauchamp 1998). In part this may be because foraging itself is a complex trait that is the outcome of multiple functional variables, and individuals differ in how these combine to determine their overall foraging strategy (Nachappa et al. 2010). For instance, some individuals are simply more active foragers than others (DeLong et al. 2021), and for any given foraging effort some will spend more time as producers and others as scroungers (Morand-Ferron et al. 2007). The interplay between sociality and individual variation in multiple components of foraging is not well understood (Morand-Ferron et al. 2011; Aplin and Morand-Ferron 2017), but critical for a full understanding of the factors that determine foraging success in natural populations.

Understanding the effects of sociality on foraging is further complicated because not all individuals in the same social group experience the same social environment. Animal social network analyses have shown that some individuals have stronger connections, are more central in the network or experience specific types of connections more often than others (Whitehead 2008; Krause et al. 2015). The temporal stability of social networks can also affect individual network position (Ansmann et al. 2012; Murphy et al. 2020). Social network position in turn affects individual foraging behaviors (Giraldeau and Caraco 2000). For instance, individuals that are more central in the network may experience greater levels of foraging competition but also have more opportunities to scrounge from others (Aplin and Morand-Ferron 2017; McMahon et al. 2024). Network density can also affect competition levels, although the direction of this effect is highly variable (Beauchamp 1998; Vásquez and Kacelnik 2000; Grueter et al. 2018; Sheppard et al. 2018). Individuals with close social ties may reduce their own foraging effort in order to maintain those ties (Firth et al. 2015). And individuals that are closely connected to others that act as sources of information—for example, individuals that discover a new food patch or innovate a new foraging technique—may benefit from those connections by being an early recipient of social information (Slagsvold and Wiebe 2011; Aplin et al. 2012; Kulahci et al. 2016; Schakner et al. 2017; McMahon et al. 2024). Likewise, the overall structure of the social network may impact on individual foraging behavior. For instance, networks that are more temporally stable (i.e. in which connections between individuals remain relatively consistent over time) may allow for more opportunities for social learning (Kulahci and Quinn 2019), but may also affect an individual’s competitiveness through the establishment of dominance hierarchies (McDonald and Shizuka 2013).

Different components of foraging success may be differentially impacted by social network structure and position. First, foraging traits differ in how much they rely on or are affected by social interactions. For example, scrounging requires the presence of other individuals, and is impacted by both social network structure and social stability (Firth et al. 2015; Aplin and Morand-Ferron 2017). In contrast to scrounging, other components of foraging behavior do not require social contact and may or may not be affected by interactions with others in a social group. For instance, individual foraging rates could be a product of individual hunger levels and extraction efficiency, but they can also be affected by social group characteristics such as group size, for example through effects of competition (Rieucau and Giraldeau 2009a) and the perception of predation risk driven by shared vigilance or the dilution effect (Stevens et al. 2007). Foraging rates can also be improved by the strength of social interactions, which may improve an individual’s “sense of security” (Voelkl et al. 2016; Heathcote et al. 2017; Mady and Blumstein 2017). Learning about beneficial sources of food and how these can be obtained is another important component of foraging that is not necessarily a social process although it is often modulated by social interactions. Second, the role of cognition, and the associated social environment effects on cognitive performance, varies across foraging traits. Learning about where food can be found and how it can be extracted is a common role for cognition in foraging, and is affected in multiple ways by the social environment, not only via social learning but also through indirect effects of competition, dominance and local enhancement (Pravosudov et al. 2003; Morand-Ferron and Quinn 2011; Avarguès-Weber and Chittka 2014; Ashton et al. 2018, 2019; Langley et al. 2018), although there is variation in these effects (Heinen et al. 2021a; Lambert and Guillette 2021). Learning is involved not only in production but also scrounging (Reichert et al. 2021), but learning should have limited effects on foraging performance when feeding on a known, readily accessible food source. The effects of sociality on learning may be especially important when environmental conditions change, and previously learned contingencies are no longer effective for finding food. When this happens, strong connections to other individuals may be beneficial when there is social learning, such that when any individual accesses a new food source others can quickly do the same (Aplin et al. 2012; Jones et al. 2017). However, strong connections could also be costly when conditions change if social information is outdated or if foraging opportunities available to some individuals are not available for others they are connected to (Beauchamp et al. 1997; Kendal et al. 2005; Hall and Kramer 2008; Rieucau and Giraldeau 2009b; Firth et al. 2016). In this case, individuals would be better off using their own private information to learn about new food sources, but some may do so more readily than others (Thornton and Malapert 2009; Kurvers et al. 2010; Pitera et al. 2025).

We tested social effects on foraging in wild mixed-species flocks of great tits (Parus major) and blue tits (Cyanistes caeruleus) in the context of an experiment in which birds had to learn which feeder in an array they could access to obtain food (Reichert et al. 2020). First, we tested whether an experimental manipulation of social stability would affect three measures of foraging success: reversal learning of a rewarded feeder location, propensity to scrounge, and overall foraging rate. On the one hand, social instability could lead to disruptions in social relationships and opportunities for social learning, resulting in slower reversal learning, and an increased tendency to scrounge. On the other hand, social instability could facilitate reversal learning through disruptions to competitive hierarchies at feeders while reducing scrounging opportunities. For either scenario, the effects of social instability should increase with each successive reversal. To our knowledge, no previous study in a wild species has manipulated the structure of the social network to determine the effects on multiple components of foraging success. Second, we tested the relationship between two metrics of individual social network positioning and foraging. The individual metrics used were the average size of the flocks birds were in when visiting feeders and weighted degree, a measure of the strength of social connections with other individuals in the network. Both of these metrics capture aspects of sociality that are likely to have multiple impacts on foraging by influencing shared vigilance, levels of competition, opportunities for social learning and expression of alternative foraging tactics (Dias 2006; Cantor et al. 2020). We examined these relationships across three different learning tasks, an initial production learning and two reversals. We predicted that the effects of sociality on foraging would vary across these learning tasks because they involve different cognitive traits that have been shown differ in how they interact with the social environment (Lambert and Guillette 2021): learning an initial association versus flexibility in updating behavior in response to environmental change (Izquierdo et al. 2017).

## MATERIALS AND METHODS

### Study system

In winter, great tits and blue tits form flocks along with several other species and move about woodland areas to forage (Ekman 1989). These flocks have a fission-fusion structure, in which individuals frequently leave and rejoin the group, resulting in variation in the social bonds among individuals (Aplin et al. 2015; Farine et al. 2015b). Multiple aspects of both their foraging behavior and their social networks can be readily observed by tagging individuals with passive integrated transponders and then monitoring their visits to RFID-equipped feeders (Firth and Sheldon 2015; Milligan et al. 2017; Cauchoix et al. 2022). Furthermore, by programming feeders to allow access to the food inside for only certain individuals, it is possible to both measure different components of foraging performance and to manipulate the social structure (Firth and Sheldon 2015). Social structure and social network positioning have important consequences for foraging performance (Aplin et al. 2012; Firth et al. 2015; Beck et al. 2024; McMahon et al. 2024) and for other important traits in these species (Farine and Sheldon 2015; Firth and Sheldon 2016). Furthermore, these social network effects are not limited to conspecifics because heterospecific social interactions also impact foraging (Aplin et al. 2012; Farine et al. 2012, 2014, 2015a).

We studied social behavior and foraging in the context of a discrimination learning experiment in a wild mixed species foraging flock focused on great tits and blue tits at Wytham Woods, Oxfordshire, UK. The general experimental design and methods are described in detail in our previous publications (Reichert et al. 2020, 2021; Morand-Ferron et al. 2022). Briefly, we used arrays of sunflower-seed feeders with radio frequency identification (RFID) antennas to monitor foraging of great tits and blue tits that had been fitted with a passive integrative transponder (PIT) tag (IB Technology, Aylesbury, UK). Every bird had a unique RFID code and we could therefore detect each visit by each individual to any of the feeders in the array. Each array consisted of five feeders arranged linearly and spaced 1 m apart. Four arrays were operative in different areas of the wood at any given time, and they were placed far enough apart that birds rarely visited feeders at more than one array. One set of four feeders was monitored from November-December 2017 and another set of four between January-February 2018. All procedures were approved by the Animal Welfare Body at University College Cork (HPRA license number AE19130-P017), and were in accordance with the Association for the Study of Animal Behaviour Guidelines for the Treatment of Animals in Behavioural Research and Teaching. All research was conducted under British Trust for Ornithology licenses as part of ongoing research in this population.

Prior to the learning experiments that contributed the data analyzed here, the experimental feeders were placed in the area to attract birds and allow them to become accustomed to obtaining food from them (for details see Reichert et al. 2020; Troisi et al. 2025). We initially placed two feeders 50 m apart in an ‘initial dispersed’ stage to attract birds to the area. We then removed those feeders and placed five feeders in a linear array, in the same position as they were used in the learning experiment. During this ‘open clustered’ stage, any tagged bird could feed from any feeder. During the learning experiments, the focus of this study, each bird was assigned to one of the five feeders in the array, and it could only get food from its assigned feeder. If it landed on the assigned feeder, a solenoid would activate, allowing the bird to push open a door to gain access to the seeds inside. If it landed on any of the other feeders, its visit would be recorded but the solenoid would not activate, so it would not be able to access the food. The monitoring of the visits and activation of the solenoid was controlled by a custom program on a ‘Darwin Board’ (Stickman Technologies Inc., UK). The learning experiment was carried out in three stages. The first was an initial learning stage in which each individual was randomly assigned to one of the five feeders in the array and then we monitored their visits over eight days to determine if and when they learned which feeder they were assigned to (see below). Second, in the first reversal stage, we reassigned birds to a new feeder in the array and monitored their visits for another eight-ten days. In contrast to the initial learning stage, in the reversal stage we applied two different ‘social stability treatments’ that determined how birds were reassigned to new feeders. At four of the eight sites (two in each year), we applied the ‘stable’ social treatment, in which the entire set of birds assigned to the same feeder during the initial learning stage were reassigned to the same (randomly chosen) new feeder. Thus, these birds would experience consistency in which other individuals in the flock would be feeding from the same rewarded feeder. At the other four sites, we applied the ‘unstable’ social treatment. In this treatment, each individual was randomly reassigned to a new feeder independently, and thus was not always consistently feeding from the same rewarded feeder with the same other individuals and may therefore have experienced disruptions in their social connections. Third, in the second reversal stage, we again reassigned birds to new feeders, applied the stable and unstable treatments as above, and monitored visits for eight to ten days.

In our previous analyses of this dataset, we provided an extensive description of demographic factors affecting variation in participation, likelihood of learning, and learning speed in this system (Reichert et al. 2020). We found both species- and individual-level variation in learning speed (Reichert et al. 2020), and in the propensity to scrounge (Reichert et al. 2021). In an analysis focused on great tits, we also found that the social stability treatment had lasting effects on the social network that carried over across contexts (Troisi et al. 2025). Here, we extend this work by asking whether an experimental manipulation of flock stability, and individuals’ social network positions, affected individual foraging behavior (learning, scrounging, foraging rate), and if so, which components were affected.

### Social network and foraging measures

In order to capture various aspects of foraging behavior, we calculated variables corresponding to three components of individual foraging strategies. Each of these variables was calculated separately for each individual in each of the three experimental stages. First, we characterized production learning behavior by calculating learning speed as the number of visits until an individual first met a criterion (following previous research) of 16 correct visits within a window of 20 consecutive visits (Reichert et al. 2020). Second, we characterized the propensity to scrounge as the proportion of rewarded visits (i.e. visits in which the bird obtained food, which could happen either by visiting the correct feeder or scrounging) that were scrounges. We defined scrounging as a visit by an individual to a feeder it was not assigned to (and therefore where it could not obtain food by producing) that took place 1 s or less after the departure of a bird that was assigned to that feeder. In such cases, the feeder door had not yet shut and the scrounger could obtain food from that feeder (Reichert et al. 2021). Third, we characterized foraging intake rate as a measure of the rate at which birds obtained food once they had learned the task, quantified as the number of visits to their rewarded feeder on the day after it had met the learning criterion.

We note that individuals of both great tits and blue tits foraged in the same mixed species flocks, and so the social networks were constructed and social network metrics calculated using data from all detected individuals, regardless of species. Monitoring at the study population also involves giving RFID tags to any captured marsh tit (Poecile palustris) and nuthatch (Sitta europaea), thus social networks also include these individuals when detected. We restrict our analyses to great tits and blue tits because only these two species interacted with the learning apparatus in large numbers. The number of individuals of each species included in the social network varied slightly between experimental stages (Table S1).

We built social networks by assigning birds’ visits to discrete flocking events using a Gaussian mixture model (Psorakis et al. 2012, 2015). We defined edges in the social network between each individual that was detected in a flocking event (Troisi et al. 2025). We then quantified the association strength between each possible pair of individuals using a simple ratio index calculated as the number of times both individuals were seen in the same flocking event divided by the total number of flocking events in which either (or both) individual was present (Cairns and Schwager 1987; Whitehead 2008). From these data we calculated the two social metrics used in analyses for each individual: 1) its average flock size (i.e. the mean number of individuals it was observed with in flocking events) and 2) weighted degree as the weighted sum of its association strengths with all other individuals in the network. These metrics were calculated separately for each of the three experimental stages.

### Statistical analyses

All statistical analyses were conducted in R version 4.5.0 (R Development Core Team 2024). Our dataset included 68 individual great tits and 115 individual blue tits that met the learning criterion in all three experimental stages. Other individuals that were detected during the experiments but did not meet the learning criteria were excluded, because the majority of these individuals made very few visits to the feeders (Reichert et al. 2020), and because we were interested in the effects of the social stability treatment on reversal learning performance, which could not be assessed in a meaningful way if birds did not meet the initial learning criterion prior to the reversal. We used separate statistical models for each of the three foraging strategy variables to determine how these were affected by the social stability treatment and an individual’s social network metrics. All models were run with the lme4 version 1.1-35.5 package (Bates et al. 2015). Where significant interactions were identified we used Tukey post-hoc tests to determine which groups were different from one another. For interactions involving categorical variables (experimental stage, species and social stability), we did post-hoc testing using the emmeans function from the emmeans version 1.10.4 package in R (Lenth 2022), which calculates estimated marginal means for each combination of factors. For interactions involving a continuous variable (average flock size or weighted degree), we did post-hoc testing using the emtrends function in the emmeans package, which tests for differences in slopes of the relationship between the continuous predictor and the foraging variable between the levels of the categorical variables (experimental stage, species). We checked model assumptions of normality or uniformity of residuals and non-collinearity of variables using the check_model function in the performance version 0.14.0 package (Lüdecke et al. 2021). Learning speed (natural log-transformed) was modeled as a Gaussian variable in a linear mixed model. Proportion scrounged was modeled as a binomial variable in a generalized linear mixed model. Foraging rate was modeled as a Gaussian variable in a linear mixed model. Raw data and code are available at https://figshare.com/s/8ddfb55d391138371d36.

For each of the three foraging variables, results for which we present in separate sections, we tested for the effects of our social stability manipulation and social variables. First, we tested for effects of the social stability treatment. For these models, we only included data from the two reversal learning stages, because the social stability treatment was applied after the initial learning stage. We included individual identity as a random effect and included the three-way interaction between species, experimental stage and social stability treatment as the main factors of interest, expecting that the effects of the social stability treatment could vary by species or between the two reversal stages. Specifically, for the reversal stages, we expected that relationships between social metrics and foraging should become stronger by the second reversal as the relationships become more established in the stable group and the difference with the unstable group more pronounced. Without specifying any specific a priori predictions, we expected that relationships between sociality and foraging could vary among species because of differences in factors like body size, dominance, and predation risk, all of which influence foraging behavior. We included three additional variables as fixed effects to control for factors that were identified in previous analyses to affect learning performance: whether or not the feeder was located at the edge of the array, and the amounts of time that an individual’s assigned and non-assigned feeders were inoperable due to equipment malfunction (Reichert et al. 2020). In our previous study we found that birds assigned to edge feeders learned more quickly in all three experimental stages (Reichert et al. 2020). Some feeders malfunctioned during the course of the experiment because of power loss or antenna failure and did not open for any of the birds or record any visits until they were repaired (further details and data on these occurrences are given in Reichert et al. 2020). We still include data from birds in such arrays, and our statistical approach aims to minimize these disruptions; we acknowledge that feeder malfunctions could have affected individual learning performance and are a source of noise for the other variables, which we assume was random with respect to the hypotheses tested.

Second, we tested for effects of the two social network position variables, average flock size and weighted degree. These models included data from all three experimental stages. Each model included individual identity as a random effect and two three-way interaction terms: species, experimental stage and average flock size, and species, experimental stage and weighted degree. Social stability treatment was not included as a factor in this model because it was not applied during the initial learning stage and because for these tests we were interested in the effects of the social network position itself, which would have been obscured by including a factor that affects social network position, as we demonstrated in a previous analysis of this dataset (Troisi et al. 2025). We again included additional fixed effects of whether or not the feeder was located at the edge of the array, and the amounts of time that an individual’s assigned and non-assigned feeders were inoperable due to equipment malfunction.

## RESULTS

There were 262530 records of the 183 focal individuals across the three experimental stages (great tits: 95818 visits, N = 68 individuals; blue tits: 166712 visits, N = 115 individuals). Across these stages, the average flock size was (mean ± SE) 13.74 ± 0.26 for great tits and 12.59 ± 0.21 for blue tits, and the average weighted degree was 6.65 ± 0.14 for great tits and 6.13 ± 0.12 for blue tits. Learning speed averaged 62.4 ± 6.9 and 63.8 ± 4.62 visits until the learning criterion was met for great tits and blue tits, respectively. For scrounging, great tits had 2347 total scrounging events (mean ± SE = 11.5 ± 1.8), and blue tits had 6552 scrounges (mean ± SE = 19.0 ± 1.3). Foraging rate averaged 51.9 ± 1.4 and 46.0 ± 0.9 visits on the day after the learning criterion was met for great tits and blue tits, respectively.

### Effects of sociality on learning speed

There was no significant three-way interaction between social stability treatment, experimental stage and species (Table 1). However, there was a significant interaction between social stability treatment and experimental stage (Table 1, Figure 1A, 1B). For both species, there was no difference between the unstable and stable groups in learning speeds in the first reversal, but in the second reversal, individuals in the unstable treatment learned more quickly (Table S2, Figure 1A). Learning speeds were not significantly different in the two social stability treatment groups in the initial learning stage, confirming that there were no differences prior to the treatment being applied (Table S3).

**Figure 1.**
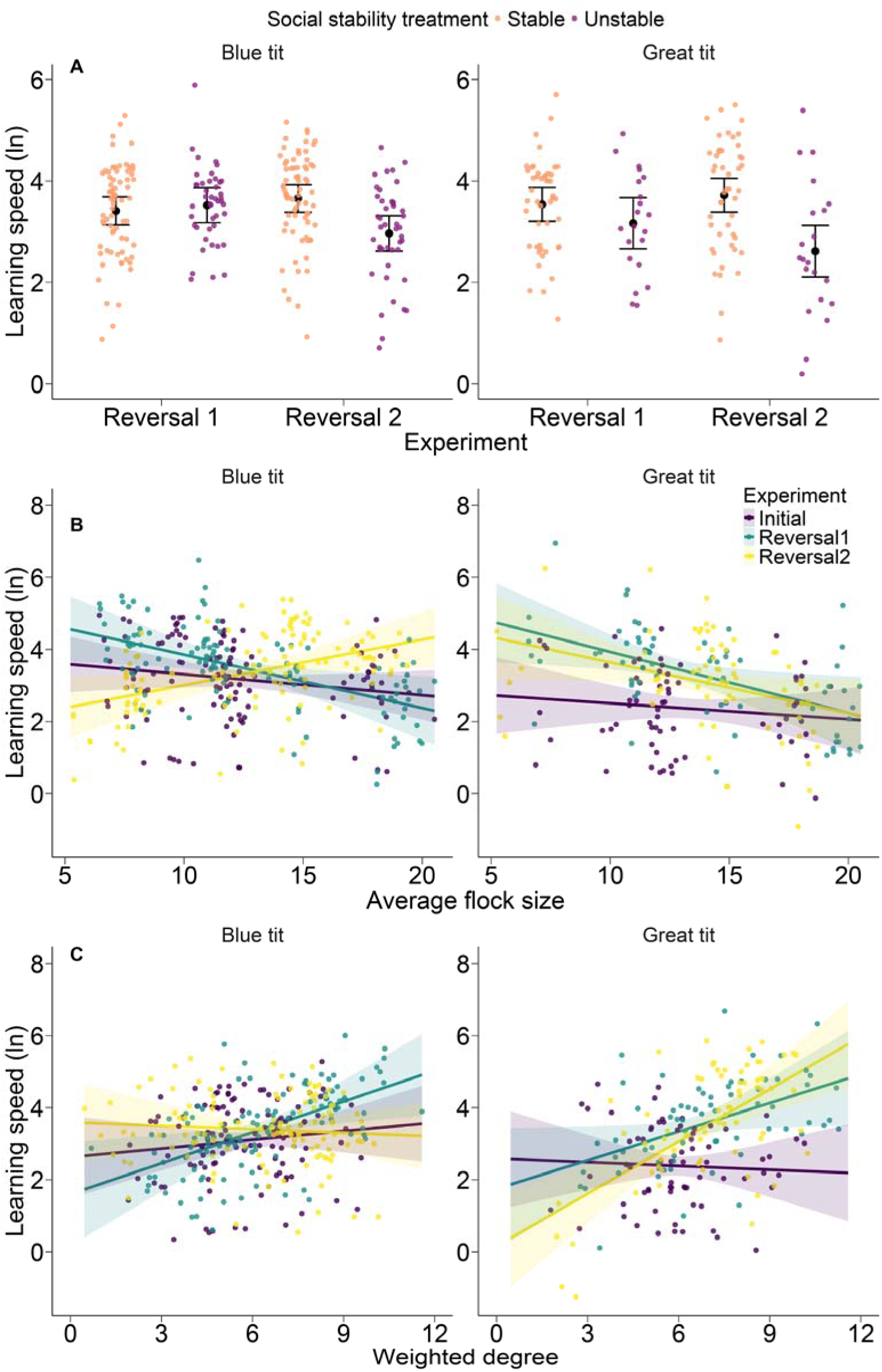
Effects of sociality on learning speed. A. Effects of social stability treatment and experimental stage on learning speed (ln-transformed) for blue tits (left) and great tits (right). Black points represent estimated marginal means (± 95% CI) from the model. Small colored points represent partial residuals from the model for each individual. B. Relationship between average flock size and learning speed for each species. Lines represent predicted estimate (± 95% CI) from the model, with separate colors for each experimental stage, and points represent partial residuals from the model for each individual. C. Relationship between weighted degree and learning speed for each species. Interpretation as in B.

**Table 1.**
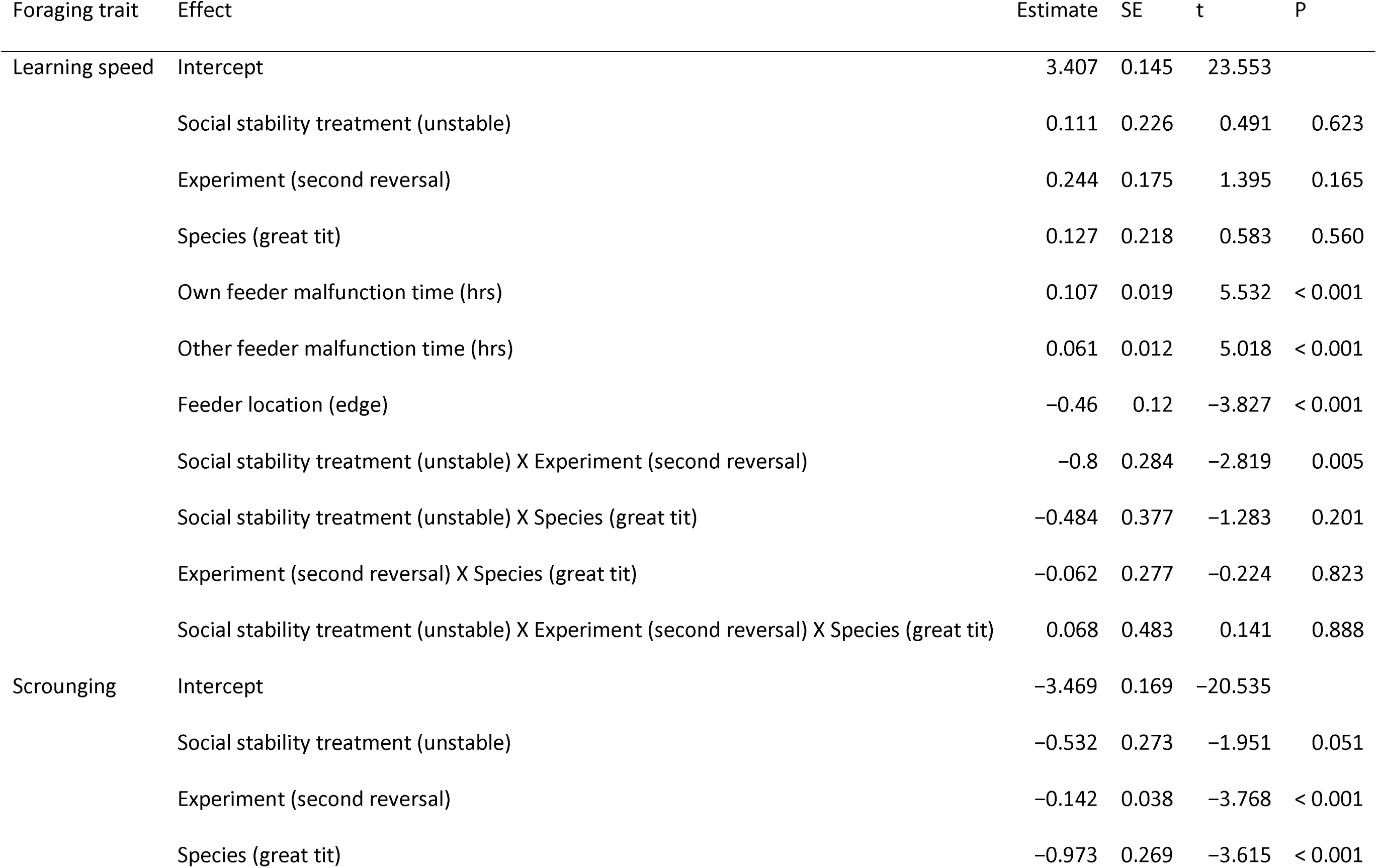
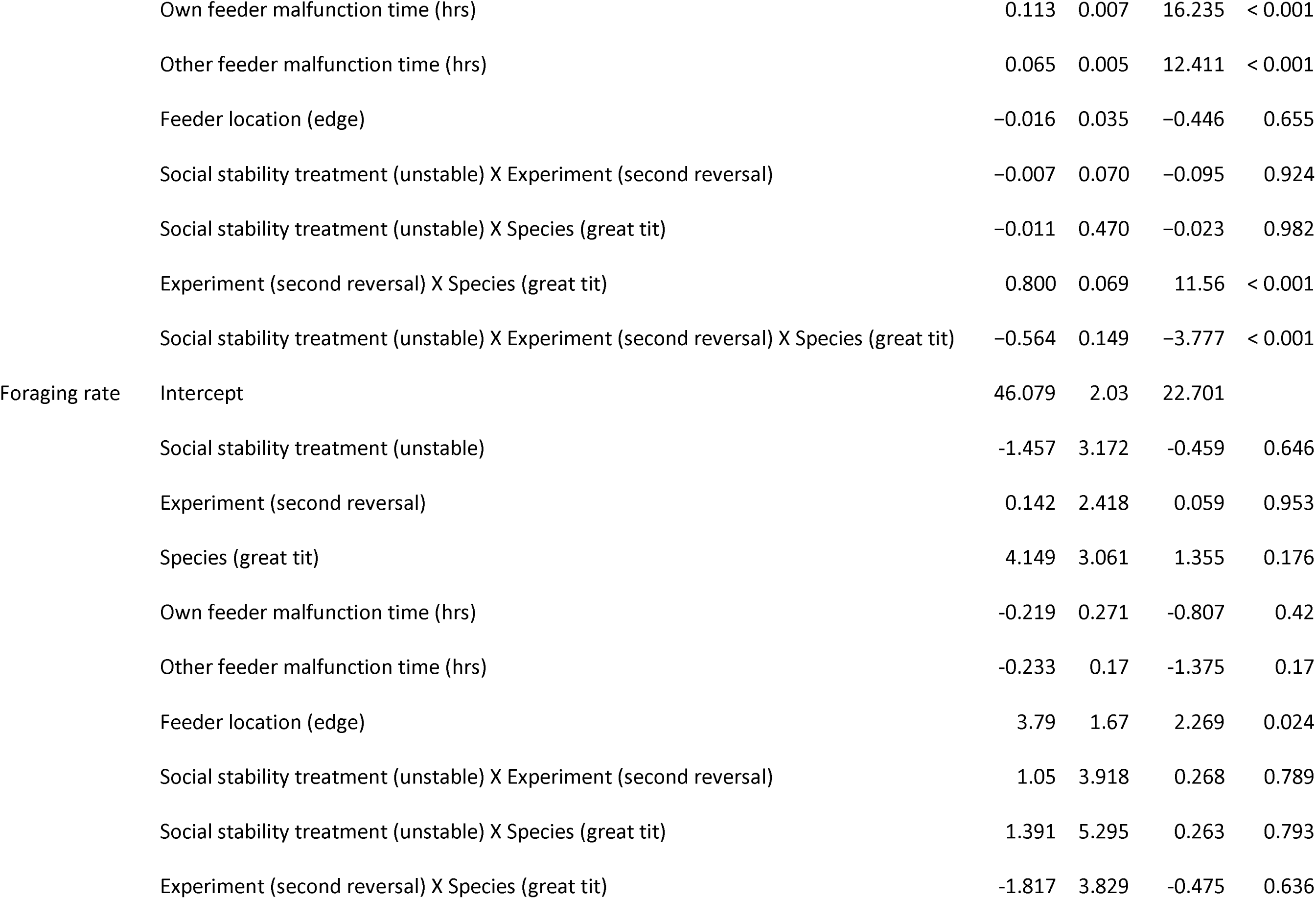
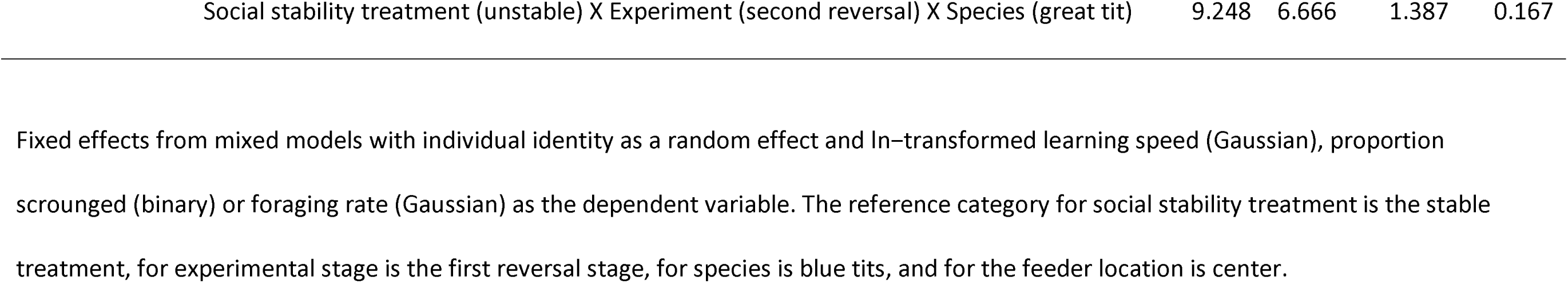
Effects of social stability on foraging traits.

| Foraging trait | Effect | Estimate | SE | t | P |
| --- | --- | --- | --- | --- | --- |
| Learning speed | Intercept | 3.407 | 0.145 | 23.553 |  |
|  | Social stability treatment (unstable) | 0.111 | 0.226 | 0.491 | 0.623 |
|  | Experiment (second reversal) | 0.244 | 0.175 | 1.395 | 0.165 |
|  | Species (great tit) | 0.127 | 0.218 | 0.583 | 0.560 |
|  | Own feeder malfunction time (hrs) | 0.107 | 0.019 | 5.532 | < 0.001 |
|  | Other feeder malfunction time (hrs) | 0.061 | 0.012 | 5.018 | < 0.001 |
|  | Feeder location (edge) | -0.46 | 0.12 | -3.827 | < 0.001 |
|  | Social stability treatment (unstable) X Experiment (second reversal) | -0.8 | 0.284 | -2.819 | 0.005 |
|  | Social stability treatment (unstable) X Species (great tit) | -0.484 | 0.377 | -1.283 | 0.201 |
|  | Experiment (second reversal) X Species (great tit) | -0.062 | 0.277 | -0.224 | 0.823 |
|  | Social stability treatment (unstable) X Experiment (second reversal) X Species (great tit) | 0.068 | 0.483 | 0.141 | 0.888 |
| Scrounging | Intercept | -3.469 | 0.169 | -20.535 |  |
|  | Social stability treatment (unstable) | -0.532 | 0.273 | -1.951 | 0.051 |
|  | Experiment (second reversal) | -0.142 | 0.038 | -3.768 | < 0.001 |
|  | Species (great tit) | -0.973 | 0.269 | -3.615 | < 0.001 |
|  | Own feeder malfunction time (hrs) | 0.113 | 0.007 | 16.235 | < 0.001 |
|  | Other feeder malfunction time (hrs) | 0.065 | 0.005 | 12.411 | < 0.001 |
|  | Feeder location (edge) | -0.016 | 0.035 | -0.446 | 0.655 |
|  | Social stability treatment (unstable) X Experiment (second reversal) | -0.007 | 0.070 | -0.095 | 0.924 |
|  | Social stability treatment (unstable) X Species (great tit) | -0.011 | 0.470 | -0.023 | 0.982 |
|  | Experiment (second reversal) X Species (great tit) | 0.800 | 0.069 | 11.56 | < 0.001 |
|  | Social stability treatment (unstable) X Experiment (second reversal) X Species (great tit) | -0.564 | 0.149 | -3.777 | < 0.001 |
| Foraging rate | Intercept | 46.079 | 2.03 | 22.701 |  |
|  | Social stability treatment (unstable) | -1.457 | 3.172 | -0.459 | 0.646 |
|  | Experiment (second reversal) | 0.142 | 2.418 | 0.059 | 0.953 |
|  | Species (great tit) | 4.149 | 3.061 | 1.355 | 0.176 |
|  | Own feeder malfunction time (hrs) | -0.219 | 0.271 | -0.807 | 0.42 |
|  | Other feeder malfunction time (hrs) | -0.233 | 0.17 | -1.375 | 0.17 |
|  | Feeder location (edge) | 3.79 | 1.67 | 2.269 | 0.024 |
|  | Social stability treatment (unstable) X Experiment (second reversal) | 1.05 | 3.918 | 0.268 | 0.789 |
|  | Social stability treatment (unstable) X Species (great tit) | 1.391 | 5.295 | 0.263 | 0.793 |
|  | Experiment (second reversal) X Species (great tit) | -1.817 | 3.829 | -0.475 | 0.636 |

|  |  |  |  |  |
| --- | --- | --- | --- | --- |
| Social stability treatment (unstable) X Experiment (second reversal) X Species (great tit) | 9.248 | 6.666 | 1.387 | 0.167 |

Fixed effects from mixed models with individual identity as a random effect and ln-transformed learning speed (Gaussian), proportion
scrounged (binary) or foraging rate (Gaussian) as the dependent variable. The reference category for social stability treatment is the stable
treatment, for experimental stage is the first reversal stage, for species is blue tits, and for the feeder location is center.

There were significant three-way interactions between species, experimental stage and both average flock size and weighted degree on learning speed (Table 2). For blue tits, individuals in larger flocks learned faster in the initial and first reversal stages, but this pattern was reversed in the second reversal stage where individuals in smaller flocks learned faster (Table S4, Figure 1B). For great tits, individuals generally learned faster in larger flocks, but, unlike in blue tits, the relationship between flock size and learning speed did not differ across experimental stages (Table S4, Figure 1B). Blue tits were relatively unaffected by weighted degree and showed no significant variation in the relationship between weighted degree and learning speed across the three experimental stages, while great tits showed a significantly different relationship between these two variables in the second reversal compared to initial learning (Table S5, Figure 1C). Specifically, great tits’ learning speeds were relatively independent of weighted degree in the initial learning stage, but by the second reversal individuals with lower weighted degrees learned more quickly than individuals with higher weighted degrees (Table S5, Figure 1C).

**Table 2.**
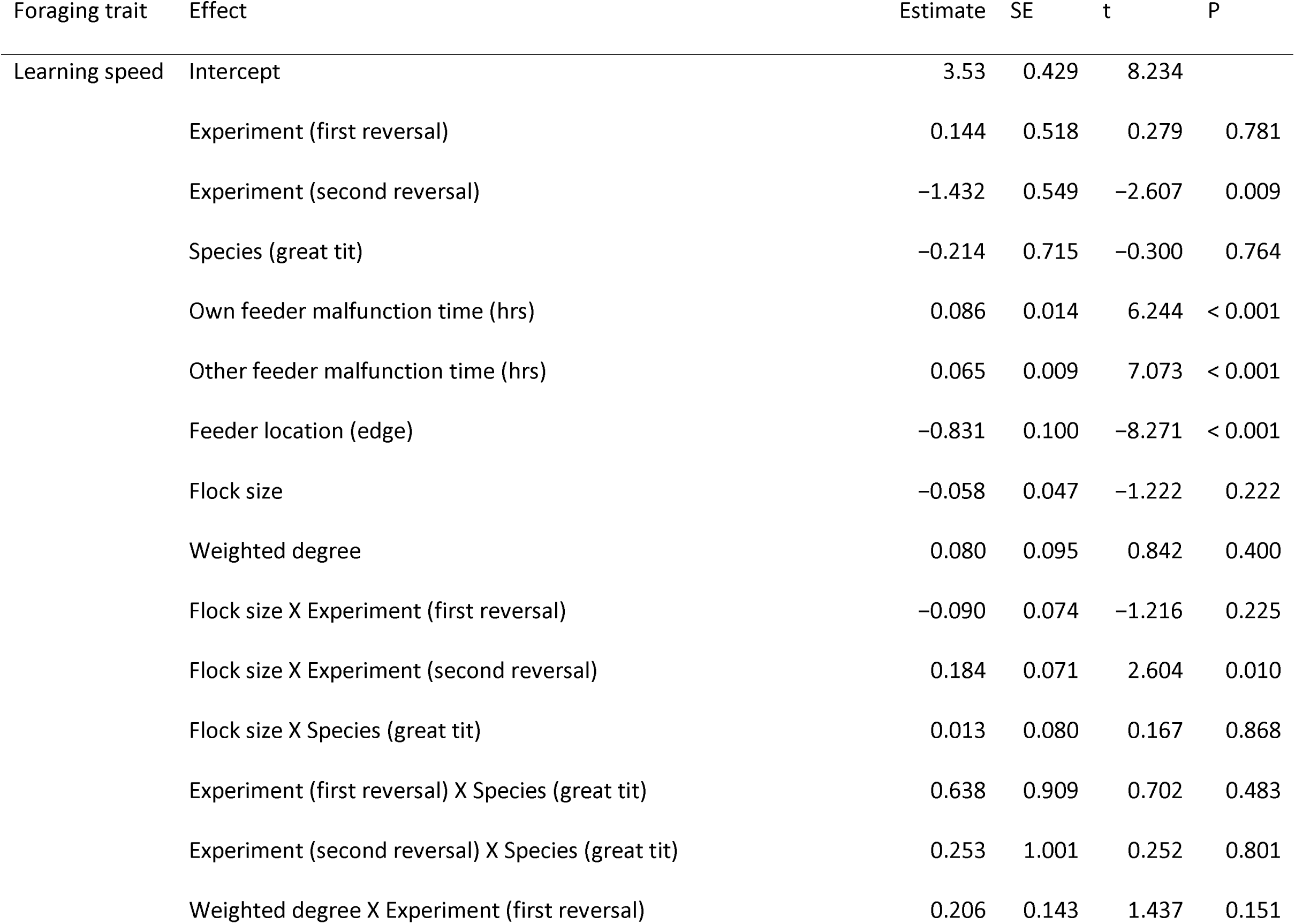
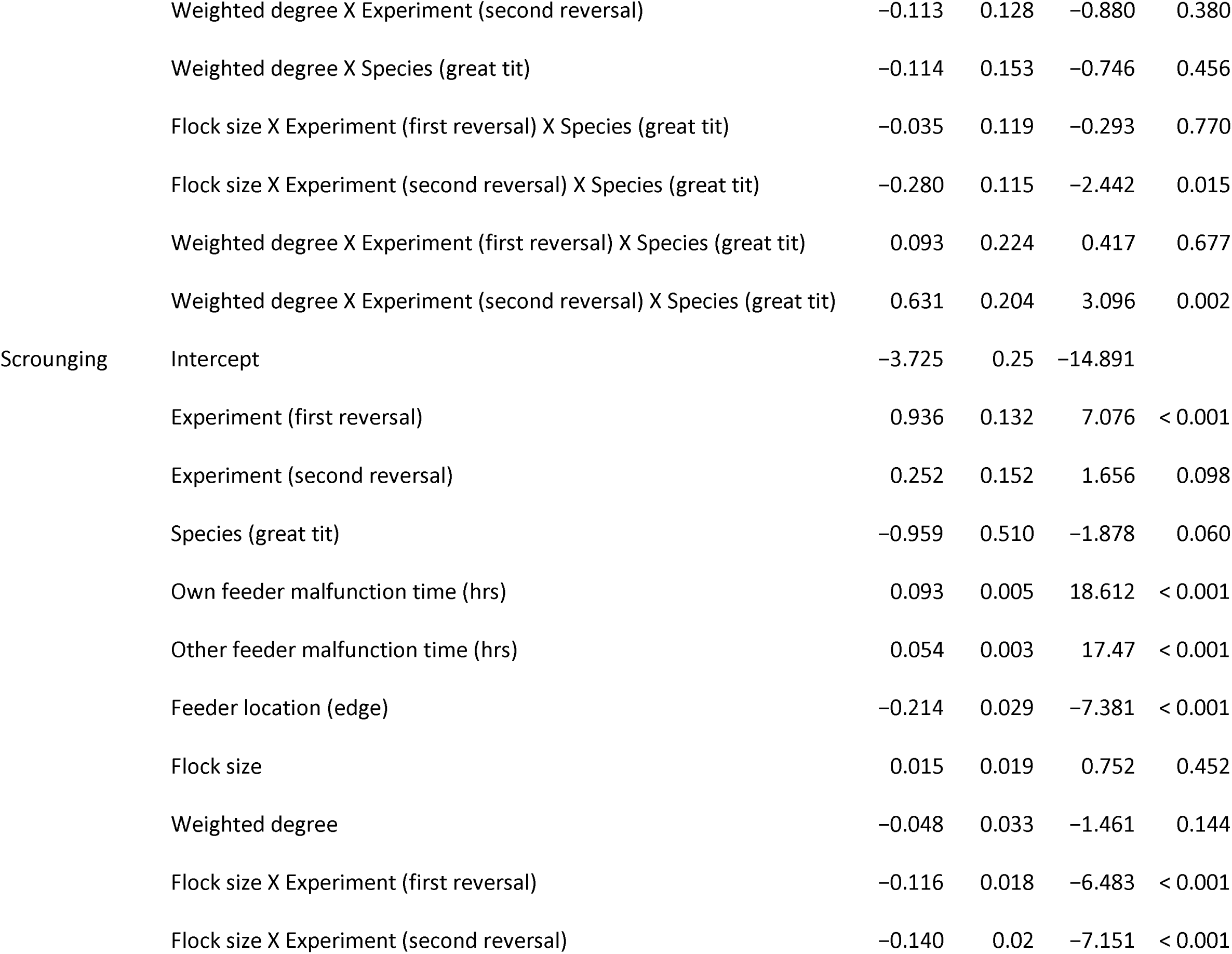
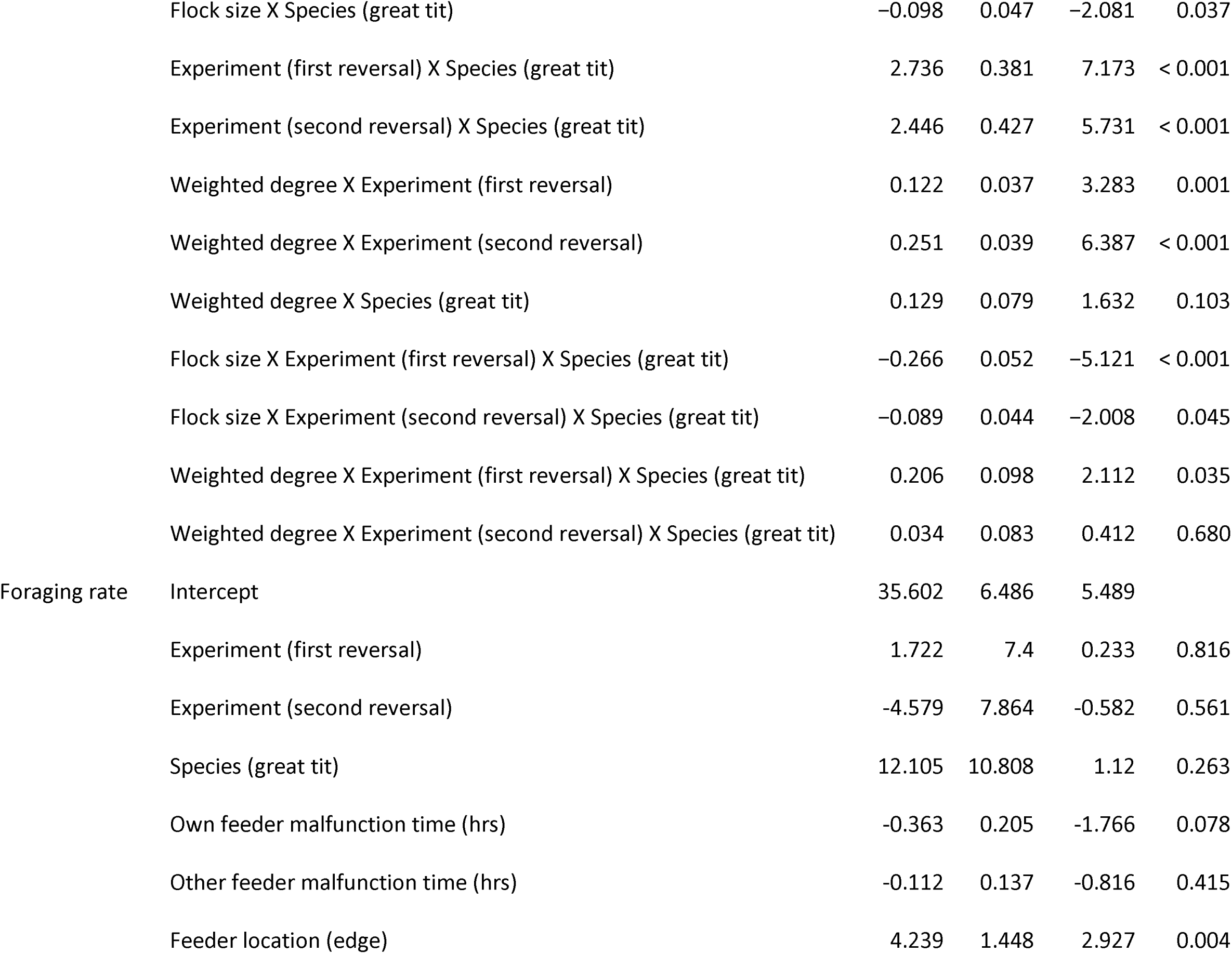
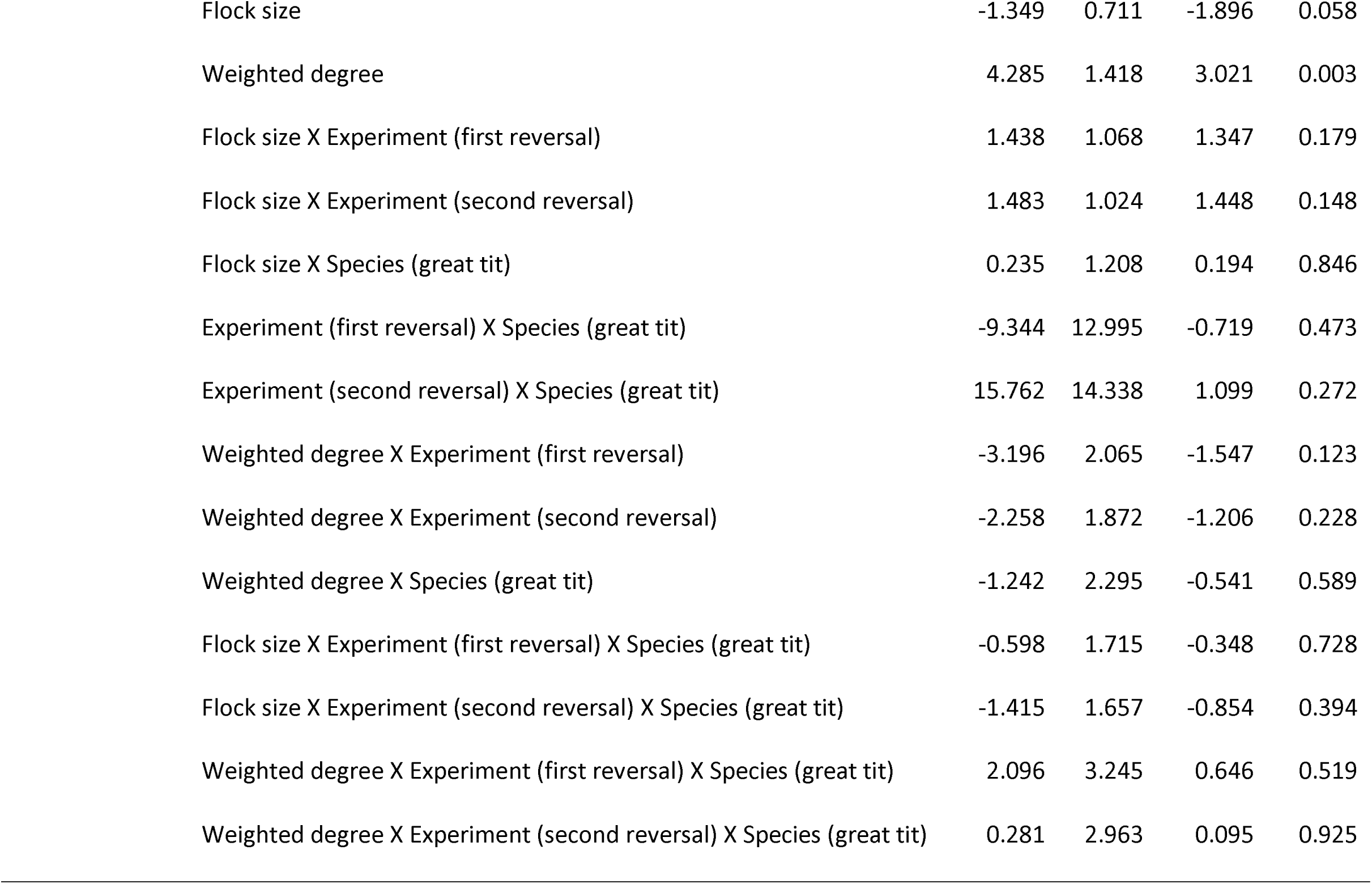
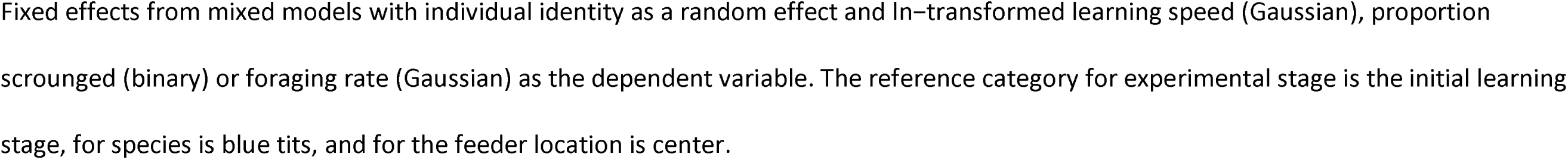
Effects of social characteristics on foraging traits.

### Effects of sociality on scrounging

There was a significant three-way interaction between species, experimental stage and social stability treatment on the proportion scrounged (Table 1). Post-hoc tests indicated that scrounging levels were significantly higher in the stable treatment compared to the unstable treatment for both species in the second reversal (Table S6, Figure 2A). However, in the first reversal there was no effect of social stability treatment on scrounging for great tits, and for blue tits there was a marginal effect, where again scrounging levels were higher in the stable compared to the unstable treatment groups (Table S6, Figure 2A). However, we noticed that in the initial learning stage, there was a main effect of social stability treatment: individuals at the sites that would be assigned to the stable treatment had higher significantly scrounging levels than individuals in the unstable treatment, indicating that there may have been a pre-existing difference between these groups for this trait, and this difference applied to both species (Table S3). However, because we found no significant effect of social stability treatment in the first reversal and then significant differences in the second reversal, the treatment had an effect over and above any differences in the populations prior to treatment.

**Figure 2.**
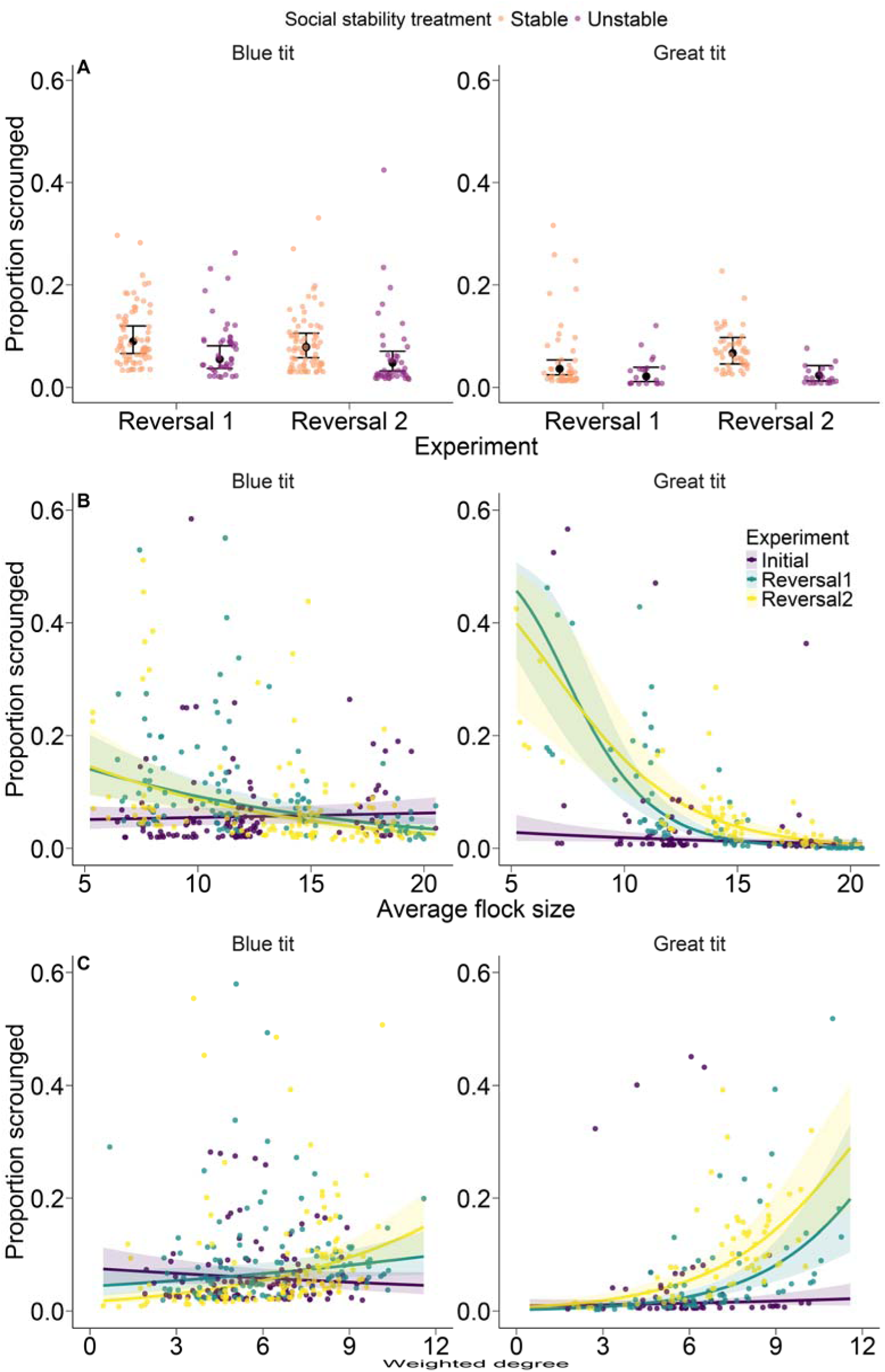
Effects of sociality on scrounging. A. Effects of social stability treatment and experimental stage on the proportion of feeds that were scrounges for blue tits (left) and great tits (right). Black points represent estimated marginal means (± 95% CI) from the model. Small colored points represent partial residuals from the model for each individual. B. Relationship between average flock size and scrounging for each species. Lines represent predicted estimate (± 95% CI) from the model, with separate colors for each experimental stage, and points represent partial residuals from the model for each individual. C. Relationship between weighted degree and scrounging for each species. Interpretation as in B. The y-axis is cut off at 0.6 for visualization purposes, there are 1, 13, and 13 outlying points not depicted in A, B, and C, respectively. Figure S1 presents the same graphs but with all data points included.

There were significant three-way interactions between species, experimental stage and both average flock size and weighted degree on proportion scrounged (Table 2). For great tits, individuals in larger flocks scrounged less often but this effect was much stronger in the reversal stages than in the initial learning stage (Table S7, Figure 2B). There was also a significant difference between the two reversal stages, with a stronger negative relationship in the first reversal than in the second reversal (Table S7, Figure 2B). For blue tits, the relationship between flock size and proportion scrounged was overall much weaker, but both reversal stages showed a more negative relationship between flock size and proportion scrounged compared to the initial learning stage (Table S7, Figure 2B). In both species there was an increasingly positive relationship between weighted degree and proportion scrounged as the experiment proceeded from the initial learning stage through the reversals (Figure 2C): individuals with higher weighted degree scrounged more often. For blue tits, the relationship became significantly stronger and more positive in comparisons between each successive experimental stage (Table S8, Figure 2C). For great tits, both reversal stages showed significantly stronger positive relationships between weighted degree and proportion scrounged compared to the initial learning stage, but there was no difference between the two reversal stages (Table S8, Figure 2C).

### Effects of sociality on foraging rate

There was no effect of the social stability treatment on foraging rate (Figure 3A). This was the case both for the full model including interactions with species and experimental stage (Table 1) and for a reduced model that did not include the interactions (Table S9). In the reduced model, there was a statistically significant effect of species, with great tits having a higher foraging rate than blue tits, and an effect of feeder location, with higher foraging rates for feeders on the edge of the array (Table S9). Foraging rates were not significantly different in the two social stability treatment groups in the initial learning stage, confirming that there were no differences prior to the treatment being applied (Table S3).

**Figure 3.**
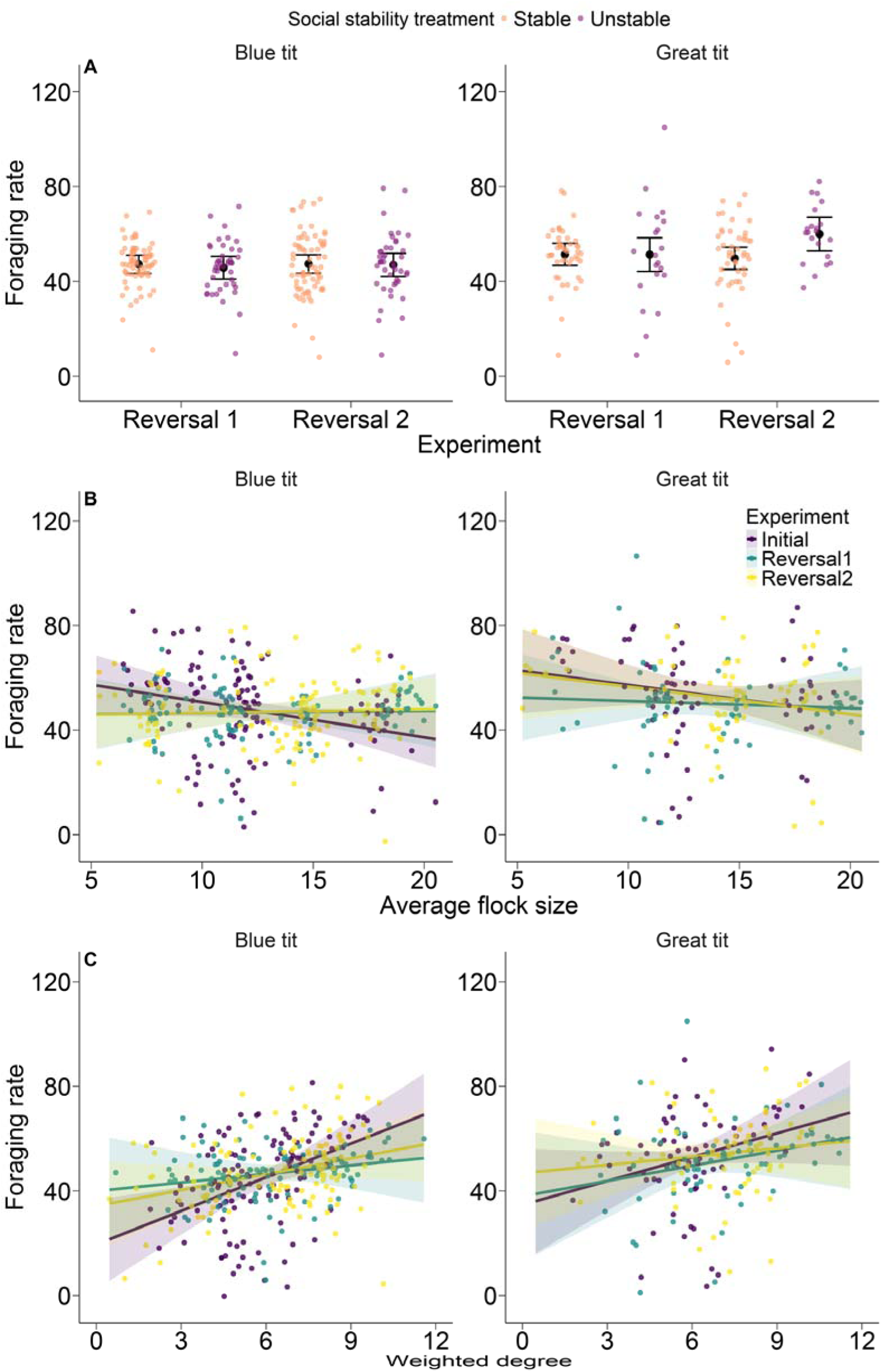
Effects of sociality on foraging rate. A. Effects of social stability treatment and experimental stage on foraging rate (number of rewarded visits on the day after the learning criterion was met) for blue tits (left) and great tits (right). Black points represent estimated marginal means (± 95% CI) from the model. Small colored points represent partial residuals from the model for each individual. B. Relationship between average flock size and foraging rate for each species. Lines represent predicted estimate (± 95% CI) from the model, with separate colors for each experimental stage, and points represent partial residuals from the model for each individual. C. Relationship between weighted degree and foraging rate for each species. Interpretation as in B.

There were no interactions between species, experimental stage and either average flock size or weighted degree (Table 2). There was, however, a main effect of weighted degree: individuals with a higher weighted degree had a higher foraging rate (Figure 3C). This effect was present in both the full model (Table 2) and in a reduced model that did not include interaction effects (Table S10). In the full model there was a marginal effect of average flock size (Table 2), although this effect was reduced and not statistically significant in the reduced model without interactions (Table S10). Foraging rates generally remained relatively constant or decreased slightly with increasing flock sizes (Figure 3B).

## DISCUSSION

Sociality had clear effects on all aspects of foraging behavior, indicating that foraging in social groups affects food intake, which could carry over to affect allocation of resources to life history traits. In a previous analysis of this dataset focused on great tits, we demonstrated that metrics of an individual’s social network position in these experiments are repeatable, indicating consistent among-individual variation in sociality (Troisi et al. 2025). Here we show that these same metrics affect foraging behavior in both great tits and blue tits, demonstrating an important consequence of individual variation in sociality and also indicating that these summary metrics of social position do indeed capture meaningful biological variation. Although there were multiple differences in effects on the two study species, these were often in the magnitude rather than the direction of the effect, which also underscores the robustness of our findings. The links among these traits were context dependent, and suggest interactions with both individual factors like variation in cognitive ability and the positive and negative effects of social grouping (reviewed in Table 3). Below, we explore and highlight the novelty in our results, and generate hypotheses for future experimental work to understand the mechanisms by which complex social structure affects different aspects of foraging success simultaneously.

**Table 3.** Summary of results and potential mechanisms.

| Social trait | Foraging trait | Effect | Potential mechanism |
| --- | --- | --- | --- |
| Social stability | Learning speed | Negative | Social instability disrupts dominance hierarchies, facilitating learning |
|  | Scrounging | Positive | More competition in stable groups, or trade-off with learning speed |
|  | Foraging rate | None | Individual variation in hunger, motivation or competitive ability |
| Flock size | Learning speed | Mostly positive | Learning is facilitated by shared vigilance, reduced risk of predation |
|  | Scrounging | Negative | Trade-off with learning speed |
|  | Foraging rate | None | Individual variation in hunger, motivation or competitive ability |
| Weighted degree | Learning speed | Negative | Individuals with strong social ties visit non-rewarded feeders with social partners |
|  | Scrounging | Positive | Stronger connections facilitate learning to scrounge, or trade-off with learning speed |
|  | Foraging rate | Positive | Increased sense of security and shared vigilance with strong social connections |

For each combination of social and foraging trait measured, we give the direction for the general trend in the relationship between the two traits, but note that all of these effects showed additional variation across species and experimental stages that in some cases resulted in a qualitatively different relationship. Potential mechanisms for the effect are given, but these are speculative and our experiments were not designed to isolate specific mechanisms.

### Effects of sociality on learning speed

We had predicted that reversal learning would be faster in the stable treatment because of increased opportunities for social learning, but we found no effect of treatment for the first reversal and a significant effect in the opposite direction for the second reversal. For both species, individuals in the unstable group were faster to learn in the second reversal. We note that a previous analysis on this study system focused on the dynamics of the learning process, and therefore including many other variables not analyzed here, found no effects of social stability treatment on reversal learning speed (Reichert et al. 2020). The difference between our current study and these previous results can be explained by our more focused analysis and including both experimental stages in the same model, which increases power and allows the contrast between the first and second reversal to be identified. Both species were affected similarly by the treatment, suggesting that the consequences of social stability have general effects on learning in a foraging context despite species differences in competitive ability and learning performance (Reichert et al. 2020, 2021). One explanation for our finding that learning performance was not better in stable social groups is that most individuals learned the task relatively quickly, potentially limiting the utility of social learning in this system (Reichert et al. 2020).

The somewhat improved learning performance in the unstable treatment in our study could have occurred because this treatment disrupted social competition and dominance relationships that built up among individuals attempting to access the same feeder. Our social network metrics do not capture such relationships because they are based on flock membership, but they are plausible, because high levels of competition and low dominance ranks are associated with reduced learning performance in many species (Chalmeau and Gallo 1993; Pravosudov et al. 2003; Langley et al. 2018). Similarly, there is evidence from multiple species, including great tits, that better problem solvers are poor competitors at feeders (Cole and Quinn 2012), so the unstable treatment may have enabled those better at problem solving to realize greater cognitive performance when otherwise negative hierarchical interactions with known individuals were reduced. However, other studies of learning in wild systems found limited effects of dominance and social stability (Heinen et al. 2021a, 2021b; Pitera et al. 2025), and other social processes caused by our manipulation could also be responsible.

For great tits, we found limited evidence for effects of average flock size on learning speed, but in general individuals with larger average flock sizes learned more quickly. For blue tits, individuals with larger average flock size also learned more quickly in the first reversal, but more slowly in the second reversal. In a previous study of great tits from the same population, rates of innovative problem solving also increased with flock size (Morand-Ferron and Quinn 2011). Thus, the costs of being in larger groups, such as increased competition, may be outweighed by other benefits, such as dilution of risk, in addition to other more subtle benefits including better cognitive performance, at least for some contexts and cognitive tasks. When we examined the effects of weighted degree, the strength of an individual’s social connections, we found that, at least for the second reversal, great tits with higher weighted degrees were slower to learn the task. In a previous study, great tits reduced their own foraging to stay with their mating partner (Firth et al. 2015). Particularly for the challenging second reversal, if similar behavior occurred in our study where birds were similarly restricted in feeder access, then this tendency to remain near strong social partners could have slowed down an individual’s own learning. Blue tits were less affected by this variable, perhaps because their cognitive performance is more strongly affected by general social competition to access feeders.

### Effects of sociality on scrounging

Scrounging rates were higher in the stable treatment during the second reversal stage for both species and, for blue tits, also in the first reversal. These results ran counter to our expectation that individuals in the unstable treatment would scrounge more because if individuals primarily scrounge from close social associates (King et al. 2009; Harten et al. 2018), the unstable treatment would have broken up those associations. However, we previously found that the across-stage correlation between the dyadic associations between individuals in the social network did not differ between the stable and unstable treatments (Troisi et al. 2025), so disruption of scrounging associations was probably not responsible for our findings of more scrounging in the stable treatment. The most likely explanation for higher scrounging in the stable treatment is that individuals in the unstable treatment were faster to learn to produce, and therefore spent less time using the scrounging tactic, given the general trade-off between producing and scrounging (Barnard and Sibly 1981).

For both great tits and blue tits, scrounging rates were lower for birds with larger average flock size in both reversal stages. These results were surprising both because we had expected larger groups to provide more opportunities to scrounge, and because individuals in larger groups generally learned more quickly, which should have provided more opportunity for scroungers. However, in a previous analysis of this dataset, we found that scrounging and production learning trade-off such that individuals that learn quickly scrounge less often (Reichert et al. 2021). As we argued above for the effects of the social stability treatment, the faster learning speed for birds in larger flocks therefore could have corresponded with reduced benefits of scrounging relative to producing. Thus, the social factors affecting production learning, in this case potentially beneficial effects of being in large groups, could have indirectly negatively influenced the expression of an alternative foraging tactic. This explanation could also apply to the positive relationship between scrounging and weighted degree in the reversals. Particularly for the second reversal, great tits with higher weighted degree were slower to learn, which could have induced more scrounging. However, determining the direction of causation is difficult. Indeed, an alternative explanation is that because effective scrounging is a learned behavior in this system (Reichert et al. 2021), birds with stronger connections may have had more opportunities to learn effective scrounging techniques or could have developed more stable producer-scrounger relationships. This would be consistent with a previous study of great tits in which birds with especially strong connections were found to forgo production to stay with their mating partner, from whom they obtained food by scrounging (Firth et al. 2015).

### Effects of sociality on foraging rate

Individuals’ foraging rates after learning the location of their rewarded feeder were not affected by the social stability treatment for either species. Foraging rate at feeders is a repeatable trait in this system (Crates et al. 2016). This individual consistency in behavior may have contributed to our findings that the overall amount of foraging activity remained relatively stable despite any changes that the social stability treatments caused to social relationships, overall social structure, and subsequent effects on interactions at the feeders. Our findings suggest that the effects of living in groups on individual foraging rates are not influenced by the ability to feed near specific individuals, which are unnecessary for general benefits of social foraging like shared vigilance and the dilution effect, though we acknowledge that these effects may have been limited by the relative ease with which food could be obtained post learning.

Foraging rates were also unaffected by average flock size. A previous study found a similar pattern in which overall visit lengths and inter-visit intervals were not affected by an experimental manipulation of density at the feeder (Regan et al. 2022). However, it also found that individuals with stronger social ties were more likely to visit feeders and that these effects interacted with social group density for some foraging behaviors (Regan et al. 2022), pointing to the importance of weighted degree in foraging. Indeed, we found that individuals had a higher foraging rate at our feeders when they had a higher weighted degree. This effect was somewhat stronger for the initial learning stage, perhaps because at this early stage scrounging levels were low and the social stability treatment had not yet affected social connections. One important consideration is that our measure of foraging rate only includes visits to our experimental feeders. Although feeders are important food sources, great tits have a broad diet in the winter even when supplemental food is available (Coomes et al. 2025), and this is likely to be the case for blue tits as well. Thus, individuals may vary in their tendency to visit feeders in the first place, and this may be related to the strength of their social relationships, which could explain our findings.

### Differences across stages and species

A clear outcome of our study is that effects of sociality on foraging varied across the experimental stages, an initial associative learning task and two subsequent reversal learning tasks. However, this was only the case for the two cognitively demanding foraging traits, learning speed and scrounging rate (which is cognitively demanding because individuals learn to scrounge in our system; (Reichert et al. 2021)), but not for foraging rate. This implies that the effects of sociality may be especially pronounced in cognitively challenging situations. The main difference between the experimental stages was in the learning task required of the birds, from an initial association to two successive reversals. Therefore, one explanation for variation across stages is that sociality interacted differently with the cognitive challenges associated with each task. Initial learning is driven by an associative process between a feeder and the reward, while reversal learning requires additional cognitive flexibility so that individuals inhibit their response to the previously rewarded feeder and instead make an association with the new rewarded feeder (Morand-Ferron et al. 2022). At the same time, some individuals were learning to become increasingly proficient at scrounging and there were opportunities to socially learn from other individuals visiting the feeders (Reichert et al. 2021). Some previous studies showed that the effects of the social environment on cognition depend on the cognitive task (Kogan et al. 2000; Schrijver et al. 2004), and our results may indicate that these findings extend to foraging behaviors involving different cognitive mechanisms in a wild population. Although further study is needed, our results point to an effect of sociality on the opportunity to learn alternative foraging tactics, adding to the evidence that cognition is shaped by social structure (Hoppitt and Laland 2013; Seyfarth and Cheney 2015; Kulahci and Quinn 2019).

Another potential explanation for our finding of variation in the effects of sociality across stages is that the overall structural properties of the global social network itself changed. Indeed, we previously showed in this dataset that average flock size and weighted degree both increased from the initial learning to the second reversal stages, and the increase in flock size was stronger for the stable treatment (Troisi et al. 2025). Where we found significant interactions between experimental stage and social network metrics on foraging behavior, the relationships between sociality and behavior were generally stronger during the reversal stages, indicating that the increased strength of social connections led to emergent impacts on foraging. At the same time, we found support for our prediction that the effects of the social stability treatment on foraging would be stronger in the second reversal than the first reversal, which indicates that multiple social disruptions have increasingly large effects on foraging. The current study and the analyses in Troisi et al. (2025) clearly show that the foraging and social processes were highly dynamic within groups during this experiment. We note that some of these social processes were likely already in flux prior to the initial learning stage. Our experimental design involved manipulating feeder placement and access to food to attract birds to the area and acclimate them to the experimental devices before we began the learning experiments. Social network traits and flock size varied even across these early stages, indicating that all individuals would have been experiencing some social instability before our social stability manipulation (Troisi et al. 2025). Indeed, some baseline level of social instability is the norm in these foraging flocks, which are characterized by a fission-fusion structure (Farine et al. 2015b). Our social stability manipulation therefore aimed to add additional instability to the social group, and our results indicate that this did have effects on foraging traits. Nevertheless, social instability can occur at different scales (e.g., at a fine scale of accessing food at different areas in a plot to a large scale of flock membership), and further work is needed to determine if social processes operating at different scales have different effects on foraging.

We calculated the social network based on social ties across species, which is appropriate given the strong tendency of great tits and blue tits to forage together. Although some social processes like mate selection will be more strongly affected by within-species social ties (Firth et al. 2018), social foraging tactics like scrounging and competition at feeders are likely to be affected by all network members, even heterospecifics. Indeed, we found that the social stability treatment had qualitatively, if not quantitatively, similar effects on both blue tits and great tits, and the impact of weighted degree and flock size was also broadly similar on most measures of foraging in both species. These common effects not only indicate real social ties that extend to heterospecifics but also give evidence for the effectiveness of our experimental manipulation and the robustness of our results. Our findings also raise important questions for future work, including whether social effects on foraging fundamentally differ between single-species and mixed-species flocks and whether the nature of among-species interactions plays a role.

## Conclusions

Both social network position and foraging emerge from a complex interplay of factors including competition, cognition, individual personality, levels of predation and other environmental characteristics, yet disentangling these factors in the wild has been difficult due to the lack of experiments. Our findings that sociality affects multiple aspects of foraging success in a wild system expand on our knowledge of social foraging and demonstrate the utility of social network metrics for capturing variation in key behavioral traits. The results demonstrate a series of costs (slower learning in stable flocks and when weighted degree was higher) and benefits (greater scrounging opportunity with stable and more connected flocks, and higher foraging rates in flocks with stronger relationships) associated with greater sociality. However, there is still much progress to be made in identifying the mechanisms behind these results. Many of our findings suggested a role of competition and so a key next step will be to better characterize competitive interactions, which is challenging from RFID data but increasingly possible with new monitoring technologies (Smith and Pinter-Wollman 2021). Our findings also raise questions of causation: an individual’s social position certainly influences its foraging behavior, but it may also be the case that foraging behavior affects social position (Kulahci and Quinn 2019), for example if some individuals serve as models for social learning or easy targets for scroungers (Harten et al. 2018). Expanded efforts to monitor and manipulate the social dynamics of wild foraging flocks are likely to yield exciting new insights into the factors that drive the expression of functional behaviors.

## AUTHOR CONTRIBUTIONS

Conceptualization: M.R., J.F., J.Q.; Data curation: M.R., C.T., J.F.; Formal analysis: M.R., C.T., J.F.; Funding acquisition: J.Q.; Investigation: M.R.; Methodology: M.R., J.F., J.Q.; Writing – original draft: M.R.; Writing – review & editing: M.R., C.T., J.F., J.Q.

## ACKNOWLEDGEMENTS

Sam Crofts and Keith McMahon assisted with bird ringing and fieldwork. Martin Whitaker helped design the selective feeders, and Gabrielle Davidson, James Savage and Iván de la Hera Fernández helped with their construction. Karen Cogan assisted with equipment sourcing and purchasing. Ben Sheldon provided access to the study system at Wytham Woods and funding to maintain the long-term study. This research was funded by the European Research Council under the European Union’s Horizon 2020 Programme (FP7/2007-2013)/ERC Consolidator Grant ‘EVOLECOCOG’ Project No. 617509, awarded to J.L.Q., and by a Science Foundation Ireland ERCSupport Grant 14/ERC/B3118 to J.L.Q.

## REFERENCES

1. Ansmann, I. C., G. J. Parra, B. L. Chilvers, and J. M. Lanyon. 2012. Dolphins restructure social system after reduction of commercial fisheries. Animal Behaviour 84:575–581.

2. Aplin, L. M., D. R. Farine, J. Morand-Ferron, and B. C. Sheldon. 2012. Social networks predict patch discovery in a wild population of songbirds. Proceedings of the Royal Society B: Biological Sciences 279:4199–4205.

3. Aplin, L. M., J. A. Firth, D. R. Farine, B. Voelkl, R. A. Crates, A. Culina, C. J. Garroway, et al. 2015. Consistent individual differences in the social phenotypes of wild great tits, Parus major. Animal Behaviour 108:117–127.

4. Aplin, L. M., and J. Morand-Ferron. 2017. Stable producer–scrounger dynamics in wild birds: Sociability and learning speed covary with scrounging behaviour. Proceedings of the Royal Society B: Biological Sciences 284:20162872.

5. Ashton, B. J., A. R. Ridley, E. K. Edwards, and A. Thornton. 2018. Cognitive performance is linked to group size and affects fitness in Australian magpies. Nature 554:364–367.

6. Ashton, B. J., A. Thornton, and A. R. Ridley. 2019. Larger group sizes facilitate the emergence and spread of innovations in a group-living bird. Animal Behaviour 158:1–7.

7. Avarguès-Weber, A., and L. Chittka. 2014. Local enhancement or stimulus enhancement? Bumblebee social learning results in a specific pattern of flower preference. Animal Behaviour 97:185–191.

8. Barnard, C. J., and R. M. Sibly. 1981. Producers and scroungers: A general model and its application to captive flocks of house sparrows. Animal Behaviour 29:543–550.

9. Bates, D., M. Maechler, B. M. Bolker, and S. Walker. 2015. Fitting linear mixed-effects models using lme4. Journal of Statistical Software 67:1–48.

10. Beauchamp, G. 1998. The effect of group size on mean food intake rate in birds. Biological Reviews 73:449–472.

11. Beauchamp, G. 2005. Does group foraging promote efficient exploitation of resources? Oikos 111:403–407.

12. Beauchamp, G., M. Belisle, and L.-A. Giraldeau. 1997. Influence of conspecific attraction on the spatial distribution of learning foragers in a patchy habitat. The Journal of Animal Ecology 66:671–682.

13. Beck, K. B., C. E. Regan, K. McMahon, S. Crofts, E. F. Cole, J. A. Firth, and B. C. Sheldon. 2024. Experimental manipulation of population density in a wild bird alters social structure but not patch discovery rate. Animal Behaviour 209:95–120.

14. Beltrão, P., A. C. R. Gomes, and G. C. Cardoso. 2022. Collective foraging: Experimentally increased competition decreases group performance exploiting a permanent resource. Functional Ecology 36:1796–1805.

15. Cairns, S. J., and S. J. Schwager. 1987. A comparison of association indices. Animal Behaviour 35:1454– 1469.

16. Cantor, M., L. M. Aplin, and D. R. Farine. 2020. A primer on the relationship between group size and group performance. Animal Behaviour 166:139–146.

17. Cauchoix, M., G. B. Jason, A. Biganzoli, J. Briot, V. Guiraud, N. El Ksabi, D. Lieuré, et al. 2022. The OpenFeederP: A flexible automated RFID feeder to measure interspecies and intraspecies differences in cognitive and behavioural performance in wild birds. Methods in Ecology and Evolution 13:1955–1961.

18. Chalmeau, R., and A. Gallo. 1993. Social constraints determine what is learned in the chimpanzee. Behavioural Processes 28:173–179.

19. Clark, C. W., and M. Mangel. 1986. The evolutionary advantages of group foraging. Theoretical Population Biology 30:45–75.

20. Cole, E. F., and J. L. Quinn. 2012. Personality and problem-solving performance explain competitive ability in the wild. Proceedings of the Royal Society B: Biological Sciences 279:1168–1175.

21. Coomes, J. R., J. Cuff, M. Reichert, G. L. Davidson, W. O. C. Symondson, and J. Quinn. 2025. Spatio-temporal variation in diet among age and sex cohorts of a model generalist bird species, the great tit Parus major: new insights revealed by DNA metabarcoding. Ecology and Evolution 15:e71565.

22. Courchamp, F., and D. W. Macdonald. 2001. Crucial importance of pack size in the African wild dog Lycaon pictus. Animal Conservation 4:169–174.

23. Crates, R. A., J. A. Firth, D. R. Farine, C. J. Garroway, L. R. Kidd, L. M. Aplin, R. Radersma, et al. 2016. Individual variation in winter supplementary food consumption and its consequences for reproduction in wild birds. Journal of Avian Biology 47:678–689.

24. Cresswell, W. 1994. Flocking is an effective anti-predation strategy in redshanks, Tringa totanus. Animal Behaviour 47:433–442.

25. Cresswell, W. 1997. Interference competition at low competitor densities in blackbirds Turdus merula. The Journal of Animal Ecology 66:461–471.

26. DeLong, J. P., S. F. Uiterwaal, and A. I. Dell. 2021. Trait-based variation in the foraging performance of individuals. Frontiers in Ecology and Evolution 9:649542.

27. Dias, R. I. 2006. Effects of position and flock size on vigilance and foraging behaviour of the scaled dove Columbina squammata. Behavioural Processes 73:248–252.

28. Drewe, J. A. 2010. Who infects whom? Social networks and tuberculosis transmission in wild meerkats. Proceedings of the Royal Society B: Biological Sciences 277:633–642.

29. Ekman, J. 1989. Ecology of non-breeding social systems of Parus. The Wilson Bulletin 101:263–288.

30. Farine, D. R., L. M. Aplin, C. J. Garroway, R. P. Mann, and B. C. Sheldon. 2014. Collective decision making and social interaction rules in mixed-species flocks of songbirds. Animal Behaviour 95:173–182.

31. Farine, D. R., L. M. Aplin, B. C. Sheldon, and W. Hoppitt. 2015a. Interspecific social networks promote information transmission in wild songbirds. Proceedings of the Royal Society B 282:20142804.

32. Farine, D. R., J. A. Firth, L. M. Aplin, R. A. Crates, A. Culina, C. J. Garroway, C. A. Hinde, et al. 2015b. The role of social and ecological processes in structuring animal populations: a case study from automated tracking of wild birds. Royal Society Open Science 2:150057.

33. Farine, D. R., C. J. Garroway, and B. C. Sheldon. 2012. Social network analysis of mixed-species flocks: Exploring the structure and evolution of interspecific social behaviour. Animal Behaviour 84:1271–1277.

34. Farine, D. R., and B. C. Sheldon. 2015. Selection for territory acquisition is modulated by social network structure in a wild songbird. Journal of Evolutionary Biology 28:547–556.

35. Firth, J. A., E. F. Cole, C. C. Ioannou, J. L. Quinn, L. M. Aplin, A. Culina, K. McMahon, et al. 2018. Personality shapes pair bonding in a wild bird social system. Nature Ecology & Evolution 2:1696–1699.

36. Firth, J. A., and B. C. Sheldon. 2015. Experimental manipulation of avian social structure reveals segregation is carried over across contexts. Proceedings of the Royal Society B 282:20142350.

37. Firth, J. A., E. F. Cole, C. C. Ioannou, J. L. Quinn, L. M. Aplin, A. Culina, K. McMahon, et al. 2016. Social carry-over effects underpin trans-seasonally linked structure in a wild bird population. Ecology Letters 19:1324–1332.

38. Firth, J. A., B. C. Sheldon, and D. R. Farine. 2016. Pathways of information transmission among wild songbirds follow experimentally imposed changes in social foraging structure. Biology Letters 12:20160144.

39. Firth, J. A., B. Voelkl, D. R. Farine, and B. C. Sheldon. 2015. Experimental evidence that social relationships determine individual foraging behavior. Current Biology 25:3138–3143.

40. Focardi, S., and E. Pecchioli. 2005. Social cohesion and foraging decrease with group size in fallow deer (Dama dama). Behavioral Ecology and Sociobiology 59:84–91.

41. Giraldeau, L. A., and T. Caraco. 2000. Social Foraging Theory. Princeton University Press, Princeton.

42. Godfrey, S. S., C. M. Bull, R. James, and K. Murray. 2009. Network structure and parasite transmission in a group living lizard, the gidgee skink, Egernia stokesii. Behavioral Ecology and Sociobiology 63:1045–1056.

43. Grueter, C. C., A. M. Robbins, D. Abavandimwe, V. Vecellio, F. Ndagijimana, T. S. Stoinski, and M. M. Robbins. 2018. Quadratic relationships between group size and foraging efficiency in a herbivorous primate. Scientific Reports 8:16718.

44. Hall, C. L., and D. L. Kramer. 2008. The economics of tracking a changing environment: competition and social information. Animal Behaviour 76:1609–1619.

45. Harten, L., Y. Matalon, N. Galli, H. Navon, R. Dor, and Y. Yovel. 2018. Persistent producer-scrounger relationships in bats. Science Advances 4:e1603293.

46. Heathcote, R. J. P., S. K. Darden, D. W. Franks, I. W. Ramnarine, and D. P. Croft. 2017. Fear of predation drives stable and differentiated social relationships in guppies. Scientific Reports 7:41679.

47. Heinen, V. K., L. M. Benedict, A. M. Pitera, B. R. Sonnenberg, E. S. Bridge, and V. V. Pravosudov. 2021a. Social dominance has limited effects on spatial cognition in a wild food-caching bird. Proceedings of the Royal Society B: Biological Sciences 288:20211784.

48. Heinen, V. K., A. M. Pitera, B. R. Sonnenberg, L. M. Benedict, E. S. Bridge, D. R. Farine, and V. V. Pravosudov. 2021b. Food discovery is associated with different reliance on social learning and lower cognitive flexibility across environments in a food-caching bird. Proceedings of the Royal Society B: Biological Sciences 288:20202843.

49. Hoppitt, W., and K. N. Laland. 2013. Social Learning. Princeton University Press, Princeton.

50. Izquierdo, A., J. L. Brigman, A. K. Radke, P. H. Rudebeck, and A. Holmes. 2017. The neural basis of reversal learning: An updated perspective. Neuroscience 345:12–26.

51. Jones, T. B., L. M. Aplin, I. Devost, and J. Morand-Ferron. 2017. Individual and ecological determinants of social information transmission in the wild. Animal Behaviour 129:93–101.

52. Kendal, R. L., I. Coolen, Y. van Bergen, and K. N. Laland. 2005. Trade-offs in the adaptive use of social and asocial learning. Advances in the Study of Behavior 35:333–379.

53. King, A. J., N. J. B. Isaac, and G. Cowlishaw. 2009. Ecological, social, and reproductive factors shape producer–scrounger dynamics in baboons. Behavioral Ecology 20:1039–1049.

54. Kogan, J. H., P. W. Franklandand, and A. J. Silva. 2000. Long-term memory underlying hippocampus-dependent social recognition in mice. Hippocampus 10:47–56.

55. Krause, J., R. James, D. W. Franks, and D. P. Croft, eds. 2015. Animal Social Networks. Oxford University Press, Oxford.

56. Krause, J., and G. Ruxton. 2002. Living in Groups. Oxford University Press, Oxford.

57. Kulahci, I. G., and J. L. Quinn. 2019. Dynamic relationships between information transmission and social connections. Trends in Ecology and Evolution 34:545–554.

58. Kulahci, I. G., D. I. Rubenstein, T. Bugnyar, W. Hoppitt, N. Mikus, and C. Schwab. 2016. Social networks predict selective observation and information spread in ravens. Royal Society Open Science 3:160256.

59. Kurvers, R. H. J. M., K. van Oers, B. a Nolet, R. M. Jonker, S. E. van Wieren, H. H. T. Prins, and R. C. Ydenberg. 2010. Personality predicts the use of social information. Ecology letters 13:829–837.

60. Lambert, C. T., and L. M. Guillette. 2021. The impact of environmental and social factors on learning abilities: a meta-analysis. Biological Reviews 96:2871–2889.

61. Langley, E. J. G., J. O. van Horik, M. A. Whiteside, and J. R. Madden. 2018. Group social rank is associated with performance on a spatial learning task. Royal Society Open Science 5:171475.

62. Lenth, R. 2022. emmeans: estimated marginal means, aka least-squares means version 1.8.3.

63. Lima, S. L., P. A. Zollner, and P. A. Bednekoff. 1999. Predation, scramble competition, and the vigilance group size effect in dark-eyed juncos (Junco hyemalis). Behavioral Ecology and Sociobiology 46:110–116.

64. Lüdecke, D., M. Ben-Shachar, I. Patil, P. Waggoner, and D. Makowski. 2021. performance: An R Package for Assessment, Comparison and Testing of Statistical Models. Journal of Open Source Software 6:3139.

65. MacNulty, D. R., A. Tallian, D. R. Stahler, and D. W. Smith. 2014. Influence of group size on the success of wolves hunting bison. PLoS ONE 9:e112884.

66. Mady, R. P., and D. T. Blumstein. 2017. Social security: are socially connected individuals less vigilant? Animal Behaviour 134:79–85.

67. McDonald, D. B., and D. Shizuka. 2013. Comparative transitive and temporal orderliness in dominance networks. Behavioral Ecology 24:511–520.

68. McMahon, K., N. M. Marples, L. G. Spurgin, H. M. Rowland, B. C. Sheldon, and J. A. Firth. 2024. Social network centrality predicts dietary decisions in a wild bird population. iScience 27:109581.

69. Milligan, N. D., R. Radersma, E. F. Cole, and B. C. Sheldon. 2017. To graze or gorge: consistency and flexibility of individual foraging tactics in tits. Journal of Animal Ecology 86:826–836.

70. Monk, C. T., M. Barbier, P. Romanczuk, J. R. Watson, J. Alós, S. Nakayama, D. I. Rubenstein, et al. 2018. How ecology shapes exploitation: a framework to predict the behavioural response of human and animal foragers along exploration–exploitation trade-offs. Ecology Letters 21:779–793.

71. Morand-Ferron, J., L.-A. Giraldeau, and L. Lefebvre. 2007. Wild Carib grackles play a producer scrounger game. Behavioral Ecology 18:916–921.

72. Morand-Ferron, J., and J. L. Quinn. 2011. Larger groups of passerines are more efficient problem solvers in the wild. Proceedings of the National Academy of Sciences 108:15898–15903.

73. Morand-Ferron, J., M. S. Reichert, and J. L. Quinn. 2022. Cognitive flexibility in the wild: Individual differences in reversal learning are explained primarily by proactive interference, not by sampling strategies, in two passerine bird species. Learning and Behavior 50:153–166.

74. Morand-Ferron, J., G. M. Wu, and L. A. Giraldeau. 2011. Persistent individual differences in tactic use in a producer-scrounger game are group dependent. Animal Behaviour 82:811–816.

75. Murphy, D., H. S. Mumby, and M. D. Henley. 2020. Age differences in the temporal stability of a male African elephant (Loxodonta africana) social network. Behavioral Ecology 31:21–31.

76. Nachappa, P., D. C. Margolies, J. R. Nechols, and T. J. Morgan. 2010. Response of a complex foraging phenotype to artificial selection on its component traits. Evolutionary Ecology 24:631–655.

77. Oom, S. P., J. A. Beecham, C. J. Legg, and A. J. Hester. 2004. Foraging in a complex environment: from foraging strategies to emergent spatial properties. Ecological Complexity 1:299–327.

78. Pitcher, T. J., A. E. Magurran, and I. J. Winfield. 1982. Fish in larger shoals find food faster. Behavioral Ecology and Sociobiology 10:149–151.

79. Pitera, A. M., V. K. Heinen, B. R. Sonnenberg, L. M. Benedict, E. S. Bridge, and V. V. Pravosudov. 2025. Social group membership does not facilitate spatial learning of fine-scale resource locations. Behavioral Ecology and Sociobiology 79:104.

80. Pravosudov, V. V., S. P. Mendoza, and N. S. Clayton. 2003. The relationship between dominance, corticosterone, memory, and food caching in mountain chickadees (Poecile gambeli). Hormones and Behavior 44:93–102.

81. Psorakis, I., S. J. Roberts, I. Rezek, and B. C. Sheldon. 2012. Inferring social network structure in ecological systems from spatio-temporal data streams. Journal of the Royal Society Interface 9:3055– 3066.

82. Psorakis, I., B. Voelkl, C. J. Garroway, R. Radersma, L. M. Aplin, R. A. Crates, A. Culina, et al. 2015. Inferring social structure from temporal data. Behavioral Ecology and Sociobiology 69:857–866.

83. R Development Core Team. 2024. R: A language and environment for statistical computing. R Foundation for Statistical Computing, Vienna, Austria.

84. Regan, C. E., K. B. Beck, K. Mcmahon, S. Crofts, J. A. Firth, and B. C. Sheldon. 2022. Social phenotype-dependent selection of social environment in wild great and blue titsP: an experimental study. Proceedings of the Royal Society B 289:20221602.

85. Reichert, M. S., S. J. Crofts, G. L. Davidson, J. A. Firth, I. G. Kulahci, and J. L. Quinn. 2020. Multiple factors affect discrimination learning performance, but not between-individual variation, in wild mixed-species flocks of birds. Royal Society Open Science 7:192107.

86. Reichert, M. S., J. Morand-Ferron, I. G. Kulahci, J. A. Firth, G. L. Davidson, S. J. Crofts, and J. L. Quinn. 2021. Cognition and covariance in the producer-scrounger game. Journal of Animal Ecology 90:2497– 2509.

87. Rieucau, G., and L.-A. Giraldeau. 2009a. Group size effect caused by food competition in nutmeg mannikins (Lonchura punctulata). Behavioral Ecology 20:421–425.

88. Rieucau, G., and L.-A. Giraldeau. 2009b. Persuasive companions can be wrong: the use of misleading social information in nutmeg mannikins. Behavioral Ecology 20:1217–1222.

89. Sansom, A., W. Cresswell, J. Minderman, and J. Lind. 2008. Vigilance benefits and competition costs in groups: do individual redshanks gain an overall foraging benefit? Animal Behaviour 75:1869–1875.

90. Schakner, Z. A., M. B. Petelle, M. J. Tennis, B. K. Van Der Leeuw, R. T. Stansell, and D. T. Blumstein. 2017. Social associations between California sea lions influence the use of a novel foraging ground. Royal Society Open Science 4:160820.

91. Schrijver, N. C. A., P. N. Pallier, V. J. Brown, and H. Würbel. 2004. Double dissociation of social and environmental stimulation on spatial learning and reversal learning in rats. Behavioural Brain Research 152:307–314.

92. Seyfarth, R. M., and D. L. Cheney. 2015. Social cognition. Animal Behaviour 103:191–202.

93. Sheppard, C. E., R. Inger, R. A. McDonald, S. Barker, A. L. Jackson, F. J. Thompson, E. I. K. Vitikainen, et al. 2018. Intragroup competition predicts individual foraging specialisation in a group-living mammal. Ecology Letters 21:665–673.

94. Silk, J. B. 2007. The adaptive value of sociality in mammalian groups. Philosophical Transactions of the Royal Society B: Biological Sciences 362:539–559.

95. Slagsvold, T., and K. L. Wiebe. 2011. Social learning in birds and its role in shaping a foraging niche. Philosophical transactions of the Royal Society of London. Series B, Biological sciences 366:969–977.

96. Smith, J. E., and N. Pinter-Wollman. 2021. Observing the unwatchable: Integrating automated sensing, naturalistic observations and animal social network analysis in the age of big data. Journal of Animal Ecology 90:62–75.

97. Snijders, L., S. Krause, A. N. Tump, M. Breuker, C. Ortiz, S. Rizzi, I. W. Ramnarine, et al. 2021. Causal evidence for the adaptive benefits of social foraging in the wild. Communications Biology 4:94.

98. Stevens, D. W., J. S. Brown, and R. C. Ydenberg, eds. 2007. Foraging Behavior and Ecology. University of Chicago Press, Chicago.

99. Thornton, A., and A. Malapert. 2009. Experimental evidence for social transmission of food acquisition techniques in wild meerkats. Animal Behaviour 78:255–264.

100. Troisi, C. A., J. A. Firth, S. J. Crofts, G. L. Davidson, M. S. Reichert, and J. L. Quinn. 2025. Effects of fine-scale changes in resource access and social stability on the sociality of foraging flocks of wild birds. Animal Behaviour 221:123071.

101. Vásquez, R. A., and A. Kacelnik. 2000. Foraging rate versus sociality in the starling Sturnus vulgaris. Proceedings of the Royal Society of London. Series B: Biological Sciences 267:157–164.

102. Voelkl, B., J. A. Firth, and B. C. Sheldon. 2016. Nonlethal predator effects on the turn-over of wild bird flocks. Scientific Reports 6:33476.

103. Waite, T. A., and K. L. Field. 2007. Foraging with others: Games social foragers play. Pages 331–362 in D.

104. W. Stephens, J. S. Brown, and R. C. Ydenberg, eds. Foraging. Behavior and Ecology. University of Chicago Press, Chicago.

105. Weber, N., S. P. Carter, S. R. X. Dall, R. J. Delahay, J. L. McDonald, S. Bearhop, and R. A. McDonald. 2013. Badger social networks correlate with tuberculosis infection. Current Biology 23:R915–R916.

106. Whitehead, H. 2008. Analyzing animal societies: Quantitative methods for vertebrate social analysis. The University of Chicago Press.

