## Supplementary Materials for "Social connections, group size and fine-scale manipulations of social stability shape learning, scrounging and foraging rate in mixed flocks of wild songbirds"

**Table S1**. Number of individuals used to calculate the social network

| Experimental Stage | Species | Number of individuals |
| --- | --- | --- |
| Initial | Blue tit | 152 |
|  | Great tit | 117 |
|  | Marsh tit | 32 |
|  | Nuthatch | 4 |
| First reversal | Blue tit | 155 |
|  | Great tit | 117 |
|  | Marsh tit | 32 |
|  | Nuthatch | 4 |
| Second reversal | Blue tit | 151 |
|  | Great tit | 114 |
|  | Marsh tit | 33 |
|  | Nuthatch | 4 |

Social networks were calculated from the record of individual visits to the feeder array. Four species were fitted with RFID tags and could be detected at the feeders, and all of these were included in the social network. Numbers fluctuated slightly across the experimental stage because not all individuals visited the feeders in all experimental stages. See Methods in the main text for details on the social network calculation.

**Table S2**. Contrasts for effects of social stability treatment and experimental phase on learning speed

| Experiment | Social Treatment | EMM | SE | Lower CI | Upper CI | Contrast Estimate | SE | t | P |
| --- | --- | --- | --- | --- | --- | --- | --- | --- | --- |
| First reversal | Stable | 3.47 | 0.11 | 3.25 | 3.69 | 0.13 | 0.20 | 0.67 | 0.50 |
|  | Unstable | 3.34 | 0.16 | 3.04 | 3.65 |  |  |  |  |
| Second reversal | Stable | 3.69 | 0.11 | 3.47 | 3.90 | 0.90 | 0.19 | 4.63 | < 0.001 |
|  | Unstable | 2.79 | 0.16 | 2.48 | 3.10 |  |  |  |  |

Estimated marginal means (EMM) are given for each combination of social stability treatment and experimental phase from the linear mixed model for effects on learning speed (see Table 1). The contrast estimate is for the comparison between the learning speeds in the stable and unstable social stability treatments for each experimental phase, with the P-value calculated using the Tukey adjustment for multiple comparisons. Values were calculated using the emmeans function from the emmeans version 1.10.4 package in R (Lenth, 2022). Note that results are combined across species because the interaction effect including species was not statistically significant (Table 1).

**Table S3**. Models comparing foraging traits in the social stability treatment groups during the initial learning stage

| Foraging trait | Effect | Estimate | SE | t | P |
| --- | --- | --- | --- | --- | --- |
| Learning speed (ln) | (Intercept) | 3.50 | 0.18 | 19.02 |  |
|  | Social stability treatment (unstable) | 0.22 | 0.22 | 0.98 | 0.33 |
|  | Species (great tit) | −0.81 | 0.21 | −3.78 | < 0.001 |
|  | Own feeder malfunction time (hrs) | 0.07 | 0.02 | 3.64 | < 0.001 |
|  | Other feeder malfunction time (hrs) | 0.07 | 0.01 | 5.21 | < 0.001 |
|  | Feeder location (edge) | −1.54 | 0.78 | −8.61 | < 0.001 |
|  | Social stability treatment (unstable) X Species (great tit) | 0.30 | 0.37 | 0.81 | 0.42 |
| Scrounging rate | (Intercept) | −2.33 | 0.24 | −9.73 |  |
|  | Social stability treatment (unstable) | −1.07 | 0.39 | −2.76 | 0.006 |
|  | Species (great tit) | −1.50 | 0.43 | −3.49 | < 0.001 |
|  | Own feeder malfunction time (hrs) | 0.02 | 0.03 | 0.59 | 0.55 |
|  | Other feeder malfunction time (hrs) | 0.02 | 0.02 | 1.03 | 0.31 |
|  | Feeder location (edge) | −1.31 | 0.34 | −3.83 | < 0.001 |
|  | Social stability treatment (unstable) X Species (great tit) | 0.56 | 0.93 | 0.60 | 0.55 |
| Foraging rate | (Intercept) | 44.94 | 3.30 | 13.60 |  |
|  | Social stability treatment (unstable) | 2.22 | 3.96 | 0.56 | 0.58 |
|  | Species (great tit) | 9.30 | 3.83 | 2.42 | 0.02 |
|  | Own feeder malfunction time (hrs) | −0.81 | 0.34 | −2.38 | 0.02 |
|  | Other feeder malfunction time (hrs) | −0.40 | 0.24 | −1.65 | 0.10 |
|  | Feeder location (edge) | 3.78 | 3.21 | 1.18 | 0.24 |
|  | Social stability treatment (unstable) X Species (great tit) | −5.97 | 6.63 | −0.90 | 0.37 |

These models were run because the social stability treatment was only applied at the first reversal stage but it is possible that there were pre-existing differences between the treatment groups, which we could test by examining these variables in the initial learning experimental stage. Interpretation as in Table 1 in the main text. Fixed effects from a linear model with the foraging trait of interest as the dependent variable (for proportion scrounged, a general linear model with the dependent variable fit with the quasibinomial family to deal with overdispersion). The reference category for social stability treatment is the stable treatment, and for species is blue tits, and for the feeder location is center.

**Table S4**. Contrasts for effects of average flock size, experimental phase and species on learning speed

| Species | Experiment | Slope | SE | Lower CI | Upper CI | Contrast | Estimate | SE | t | P |
| --- | --- | --- | --- | --- | --- | --- | --- | --- | --- | --- |
| Blue tit | Initial | −0.06 | 0.05 | −0.15 | 0.04 | Initial − First reversal | 0.09 | 0.07 | 1.22 | 0.44 |
|  | First reversal | −0.15 | 0.06 | −0.27 | −0.03 | Initial − Second reversal | −0.18 | 0.07 | −2.60 | 0.03 |
|  | Second reversal | 0.13 | 0.05 | 0.02 | 0.23 | First reversal − Second reversal | −0.27 | 0.08 | −3.47 | 0.002 |
| Great tit | Initial | −0.04 | 0.06 | −0.17 | 0.08 | Initial − First reversal | 0.12 | 0.09 | 1.35 | 0.37 |
|  | First reversal | −0.17 | 0.07 | −0.31 | −0.03 | Initial − Second reversal | 0.10 | 0.09 | 1.07 | 0.53 |
|  | Second reversal | −0.14 | 0.07 | −0.28 | −0.004 | First reversal − Second reversal | −0.03 | 0.09 | −0.30 | 0.96 |

Estimated marginal trends are given for the relationship between average flock size and learning speed for each experimental phase and species, from the linear mixed model for effects on learning speed (see Table 2). The contrast column on the right side of the table compares slopes for each combination of experimental phases, along with P-values testing the statistical difference between slopes calculated using the Tukey adjustment for multiple comparisons. Values were calculated using the emtrends function from the emmeans version 1.10.4 package in R (Lenth, 2022).

**Table S5**. Contrasts for effects of weighted degree, experimental phase and species on learning speed

| Species | Experiment | Slope | SE | Lower CI | Upper CI | Contrast | Estimate | SE | t | P |
| --- | --- | --- | --- | --- | --- | --- | --- | --- | --- | --- |
| Blue tit | Initial | 0.08 | 0.09 | −0.11 | 0.27 | Initial − First reversal | −0.21 | 0.14 | −1.44 | 0.32 |
|  | First reversal | 0.29 | 0.11 | 0.07 | 0.51 | Initial − Second reversal | 0.11 | 0.13 | 0.88 | 0.65 |
|  | Second reversal | −0.03 | 0.09 | −0.21 | 0.14 | First reversal − Second reversal | 0.32 | 0.14 | 2.27 | 0.06 |
| Great tit | Initial | −0.03 | 0.12 | −0.27 | 0.20 | Initial − First reversal | −0.30 | 0.17 | −1.73 | 0.20 |
|  | First reversal | 0.26 | 0.13 | 0.01 | 0.52 | Initial − Second reversal | −0.52 | 0.16 | −3.27 | 0.003 |
|  | Second reversal | 0.48 | 0.11 | 0.26 | 0.70 | First reversal − Second reversal | −0.22 | 0.17 | −1.31 | 0.39 |

Estimated marginal trends are given for the relationship between weighted degree and learning speed for each experimental phase and species, from the linear mixed model for effects on learning speed (see Table 2). The contrast column on the right side of the table compares slopes for each combination of experimental phases, along with P-values testing the statistical difference between slopes calculated using the Tukey adjustment for multiple comparisons. Values were calculated using the emtrends function from the emmeans version 1.10.4 package in R (Lenth, 2022).

**Table S6**. Contrasts for effects of social stability treatment and experimental phase on proportion scrounged

| Species | Experiment | Social Treatment | EMM | SE | Lower CI | Upper CI | Contrast Estimate | SE | t | P |
| --- | --- | --- | --- | --- | --- | --- | --- | --- | --- | --- |
| Blue tit | First reversal | Stable | −3.23 | 0.17 | −3.56 | −2.90 | 0.53 | 0.27 | 1.95 | 0.051 |
|  |  | Unstable | −3.76 | 0.21 | −4.18 | −3.34 |  |  |  |  |
|  | Second reversal | Stable | −3.37 | 0.17 | −3.70 | −3.04 | 0.54 | 0.27 | 1.98 | 0.048 |
|  |  | Unstable | −3.91 | 0.21 | −4.33 | −3.49 |  |  |  |  |
| Great tit | First reversal | Stable | −4.20 | 0.21 | −4.62 | −3.79 | 0.54 | 0.38 | 1.41 | 0.16 |
|  |  | Unstable | −4.75 | 0.32 | −5.38 | −4.11 |  |  |  |  |
|  | Second reversal | Stable | −3.55 | 0.21 | −3.95 | −3.14 | 1.11 | 0.38 | 2.93 | 0.003 |
|  |  | Unstable | −4.66 | 0.32 | −5.28 | −4.04 |  |  |  |  |

Estimated marginal means are given for each combination of social stability treatment, experimental phase and species, from the generalized linear mixed model for effects on proportion scrounged (see Table 1). The contrast estimate is for the comparison between the proportion scrounged in the stable and unstable social stability treatments for each experimental phase, with the P-value calculated using the Tukey adjustment for multiple comparisons. Values were calculated using the emmeans function from the emmeans version 1.10.4 package in R (Lenth, 2022).

**Table S7**. Contrasts for effects of average flock size and experimental phase on proportion scrounged

| Species | Experiment | Slope | SE | Lower CI | Upper CI | Contrast | Estimate | SE | t | P |
| --- | --- | --- | --- | --- | --- | --- | --- | --- | --- | --- |
| Blue tit | Initial | 0.01 | 0.02 | −0.02 | 0.05 | Initial − First reversal | 0.12 | 0.02 | 6.48 | < 0.001 |
|  | First reversal | −0.10 | 0.02 | −0.15 | −0.05 | Initial − Second reversal | 0.14 | 0.02 | 7.15 | < 0.001 |
|  | Second reversal | −0.13 | 0.03 | −0.18 | −0.07 | First reversal − Second reversal | 0.02 | 0.02 | 1.23 | 0.43 |
| Great tit | Initial | −0.08 | 0.04 | −0.17 | −0.0002 | Initial − First reversal | 0.38 | 0.05 | 7.95 | < 0.001 |
|  | First reversal | −0.47 | 0.05 | −0.56 | −0.37 | Initial − Second reversal | 0.23 | 0.04 | 5.86 | < 0.001 |
|  | Second reversal | −0.31 | 0.05 | −0.41 | −0.22 | First reversal − Second reversal | −0.15 | 0.05 | −3.36 | 0.002 |

Estimated marginal trends are given for the relationship between average flock size and proportion scrounged for each experimental phase and species, from the generalized linear mixed model for effects on proportion scrounged (see Table 2). The contrast column on the right side of the table compares slopes for each combination of experimental phases, along with P-values testing the statistical difference between slopes calculated using the Tukey adjustment for multiple comparisons. Values were calculated using the emtrends function from the emmeans version 1.10.4 package in R (Lenth, 2022).

**Table S8**. Contrasts for effects of weighted degree and experimental phase on proportion scrounged

| Species | Experiment | Slope | SE | Lower CI | Upper CI | Contrast | Estimate | SE | t | P |
| --- | --- | --- | --- | --- | --- | --- | --- | --- | --- | --- |
| Blue tit | Initial | −0.05 | 0.03 | −0.11 | 0.02 | Initial − First reversal | −0.12 | 0.04 | −3.28 | 0.003 |
|  | First reversal | 0.07 | 0.04 | 0.002 | 0.15 | Initial − Second reversal | −0.25 | 0.04 | −6.39 | < 0.001 |
|  | Second reversal | 0.20 | 0.04 | 0.13 | 0.27 | First reversal − Second reversal | −0.13 | 0.04 | −3.64 | < 0.001 |
| Great tit | Initial | 0.08 | 0.07 | −0.06 | 0.22 | Initial − First reversal | −0.33 | 0.09 | −3.67 | < 0.001 |
|  | First reversal | 0.41 | 0.08 | 0.26 | 0.56 | Initial − Second reversal | −0.28 | 0.07 | −3.97 | < 0.001 |
|  | Second reversal | 0.37 | 0.05 | 0.26 | 0.47 | First reversal − Second reversal | 0.04 | 0.07 | 0.62 | 0.81 |

Estimated marginal trends are given for the relationship between weighted degree and proportion scrounged for each experimental phase and species, from the generalized linear mixed model for effects on proportion scrounged (see Table 2). The contrast column on the right side of the table compares slopes for each combination of experimental phases, along with P-values testing the statistical difference between slopes calculated using the Tukey adjustment for multiple comparisons. Values were calculated using the emtrends function from the emmeans version 1.10.4 package in R (Lenth, 2022).

**Table S9**. Effects of social stability on foraging rate from a model with no interaction terms.

| Effect | Estimate | SE | t | P |
| --- | --- | --- | --- | --- |
| Intercept | 44.926 | 1.728 | 25.994 |  |
| Social stability treatment (unstable) | 1.175 | 2.07 | 0.568 | 0.571 |
| Experiment (second reversal) | 0.888 | 1.513 | 0.587 | 0.558 |
| Species (great tit) | 5.251 | 1.963 | 2.675 | 0.008 |
| Own feeder malfunction time (hrs) | −0.137 | 0.269 | −0.512 | 0.609 |
| Other feeder malfunction time (hrs) | −0.268 | 0.169 | −1.586 | 0.114 |
| Feeder location (edge) | 3.755 | 1.666 | 2.254 | 0.025 |

Fixed effects from a linear mixed model with individual identity as a random effect and foraging rate as the dependent variable. This model differs from Table 1 in the main text because it does not include the interactions (three-way or any pairwise interactions) between experimental stage, social stability treatment and species. Reference categories for factors as in Table 1 in the main text.

**Table S10**. Effects of social characteristics on foraging rate from a model with no interaction terms.

| Effect | Estimate | SE | t | P |
| --- | --- | --- | --- | --- |
| Intercept | 38.091 | 3.333 | 11.427 |  |
| Experiment (first reversal) | −0.987 | 1.685 | −0.585 | 0.559 |
| Experiment (second reversal) | −0.105 | 1.683 | −0.062 | 0.95 |
| Species (great tit) | 5.376 | 1.845 | 2.914 | 0.004 |
| Flock size | −0.624 | 0.398 | −1.57 | 0.117 |
| Weighted degree | 2.446 | 0.706 | 3.466 | < 0.001 |
| Own feeder malfunction time (hrs) | −0.333 | 0.198 | −1.678 | 0.094 |
| Other feeder malfunction time (hrs) | −0.121 | 0.129 | −0.943 | 0.346 |
| Feeder location (edge) | 4.413 | 1.429 | 3.088 | 0.002 |

Fixed effects from a linear mixed model with individual identity as a random effect and foraging rate as the dependent variable. This model differs from Table 2 in the main text because it does not include the interactions (three-way or any pairwise interactions) between experimental stage, species and average flock size or weighted degree. Reference categories for factors as in Table 1 in the main text.

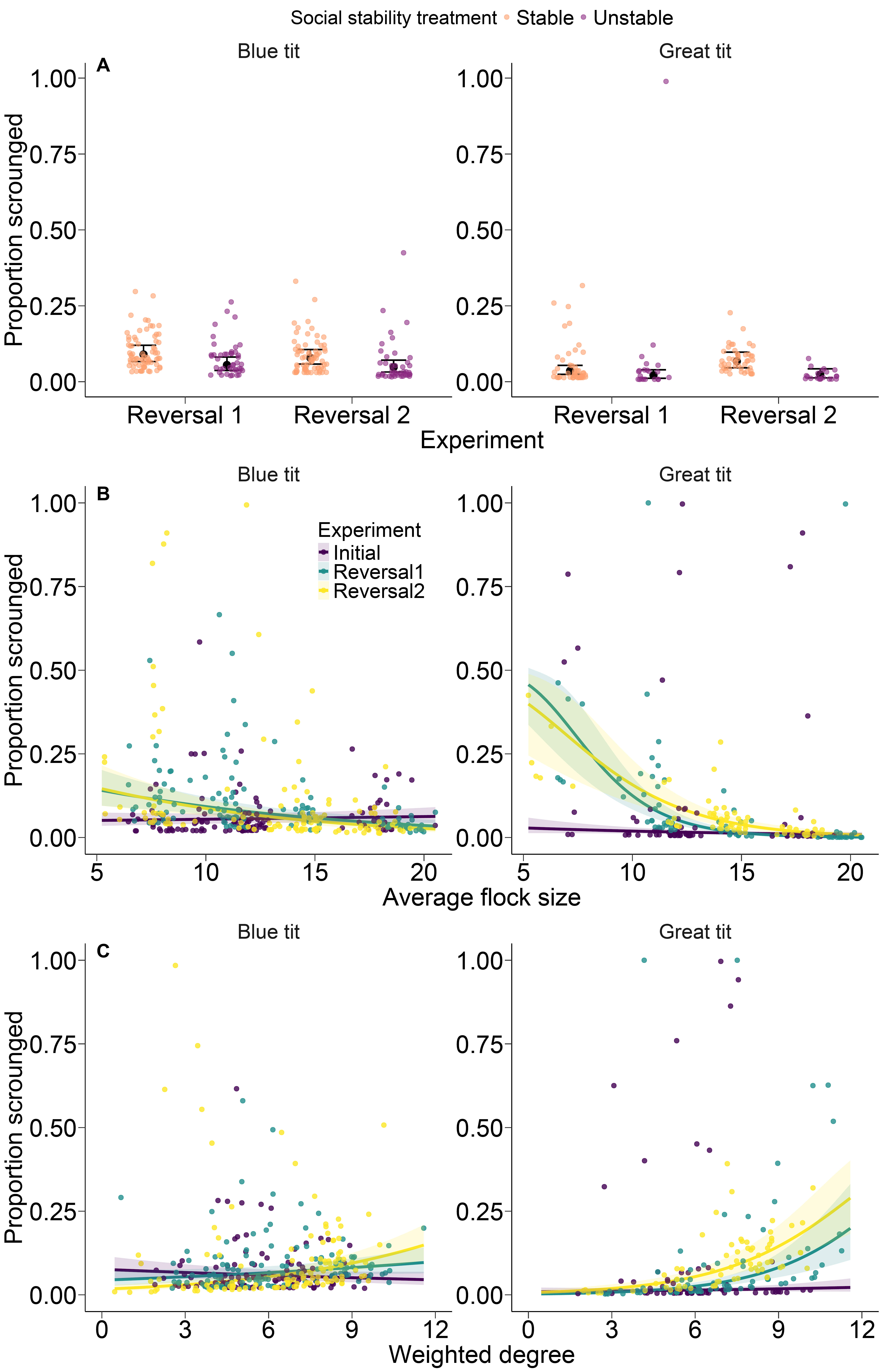

**Figure S1**. As in Figure 2 in main text, but showing all data points. The y-axis in Figure 2 was cut off at 0.6 to facilitate visualization, here the full dataset including outlying points is depicted.

References

Lenth, R. (2022). *emmeans: Estimated marginal means, aka least−squares means version 1.8.3* [Computer software].
